# Insights into energy balance dysregulation from a mouse model of methylmalonic aciduria

**DOI:** 10.1101/2021.11.30.470541

**Authors:** Marie Lucienne, Raffaele Gerlini, Birgit Rathkolb, Julia Calzada-Wack, Patrick Forny, Stephan Wueest, Andres Kaech, Florian Traversi, Merima Forny, Céline Bürer, Antonio Aguilar-Pimentel, Martin Irmler, Johannes Beckers, Sven Sauer, Stefan Kölker, Joseph P. Dewulf, Guido T. Bommer, Daniel Hoces, Valerie Gailus-Durner, Helmut Fuchs, Jan Rozman, D Sean Froese, Matthias R. Baumgartner, Martin Hrabě de Angelis

## Abstract

Inherited disorders of mitochondrial metabolism, including isolated methylmalonic aciduria (MMAuria), present unique challenges to energetic homeostasis by disrupting energy producing pathways. To better understand global responses to energy shortage, we investigated a hemizygous mouse model of methylmalonyl-CoA mutase (Mmut) type MMAuria. We found Mmut mutant mice to have reduced appetite, energy expenditure and body mass compared to littermate controls, along with a relative reduction in lean mass but increase in fat mass. Brown adipose tissue showed a process of whitening, in line with lower body surface temperature and lesser ability to cope with cold challenge. Mutant mice had dysregulated plasma glucose, delayed glucose clearance and a lesser ability to regulate energy sources when switching from the fed to fasted state, while liver investigations indicated metabolite accumulation and altered expression of peroxisome proliferator-activated receptor and Fgf21-controlled pathways. Together, these indicate hypometabolism, energetic inflexibility and increased stores at the expense of active tissue as energy shortage consequences.

## Introduction

Survival during periods of nutritional insufficiency (e.g. fasting) involves a coordinated metabolic response at the cellular and systemic levels. Adaptation in these periods requires a metabolic flexibility which may include switching of energetic sources, effective fuel management, and in more extreme cases, a reduction of metabolic rates and body temperature^1^. Inherited disorders of energy metabolism pose a unique challenge to such mechanisms, as they not only introduce chronic energy insufficiency but also interfere with adaptive mechanisms.

Isolated methylmalonic aciduria (MMAuria) is an autosomal recessive inherited inborn error of propionate metabolism caused by deficiency of the vitamin B_12_-dependant enzyme methylmalonyl-CoA mutase (MMUT). Located in the mitochondrion, MMUT catalyses the isomerization of L-methylmalonyl-CoA into succinyl-CoA, an intermediate of the tricarboxylic acid cycle, mediating an important step in catabolism of odd-chain fatty acids, cholesterol and the amino acids valine, isoleucine, methionine and threonine, and may provide up to 7-8% of daily ATP production^2^. MMAuria represents a proto-typical inherited disorder of energy metabolism in that the primary defect leads to a direct reduction of energy generation (via the tricarboxylic acid cycle), while disease-associated secondary mechanisms result in cascading disruption of other energetic pathways^3^. In the case of MMAuria, such secondary disruptions include inhibition of pyruvate dehydrogenase^4^ and citrate synthase^5^, potentially disrupting glucose oxidation, and inhibition of N-acetylglutamate synthase^6^ interfering with urea cycle function. Together, these primary and secondary disease mechanisms lead to a complex clinical picture characterized by a failure to thrive and acute crises, often triggered by a catabolic state, and by chronic progression with long-term complications in the kidney, brain and liver^7–9^.

Clinical management of patients with MMAuria is performed through pharmacological and dietary regimens that aim at keeping patients in an anabolic state, while limiting ingestion of precursor amino acids, and by replacing missing or potentially helpful molecules such as carnitine^7^. Nevertheless, many long-term complications are progressive and patients remain metabolically unstable^7,8^, suggesting that the energetic and metabolic needs of affected individuals are not fully met by these symptomatic treatments. A likely explanation is that these patients have an incomplete or mal-adaptive response to chronic energy shortage which is not currently addressed by these measures.

To understand better the metabolic basis of long-term complications in MMAuria, and how these might relate to chronic energy shortage, we have utilized a hemizygous mouse model of MMAuria, which combines a knock-in (ki) allele based on the MMUT-p.Met700Lys patient missense mutation with a knock-out (ko) allele of the same gene (*Mmut*-ko/ki)^10^. This model has the advantage of circumventing the neonatal lethality of *Mmut*-ko/ko null mutants^10,11^, and displays a strong metabolic phenotype accompanied by many clinical features of MMAuria including a pronounced failure to thrive, which are strengthened when challenged with a 51%-protein diet from day 12 of life^13^. Here, we interrogated metabolic adaptations from the whole animal to the molecular level, using body composition analysis, indirect calorimetry, blood biochemistry, histological analysis and transcriptomics. Our findings of altered fat amount and type, reduced feeding, energy expenditure and glucose clearance, and altered fibroblast growth factor 21 (Fgf21) and peroxisome proliferator-activated receptor (Ppar) expression shed light on the mechanisms and adaptations behind energy imbalance in MMAuria and provide insight into metabolic responses to chronic energy shortage.

## Results

### Body composition alteration towards less lean and more fat mass

We have previously shown that *Mmut*-ko/ki (mutant) mice on a high-protein diet are smaller than their *Mmut*-ki/wt (control) littermates and exhibit biochemical and clinical manifestations of MMAuria^13^. Here we set out to examine the metabolic basis of these disturbances by screening two cohorts of approximately 60 animals, totalling 117 mice, including females (Mmut-ki/wt, n=28; *Mmut*-ko/ki, n=31) and males (Mmut-ki/wt, n=28; *Mmut*-ko/ki, n=30) **(Fig. 1, Supplementary Fig.** 1). In line with previous findings^13^, mutant mice of both cohorts and sexes had decreased body mass compared to controls **(Fig. 1a).** Analysis of body composition by time domain-nuclear magnetic resonance suggested that mutant mice had a different proportion of lean mass **(Fig. 1b)** and fat mass **(Fig. 1c)** than controls when adjusted for body mass. Indeed, while absolute fat and lean mass values were reduced in mutant animals, females also show an unexpectedly higher proportion of fat mass compared to body mass. **(Fig. 1d).** This was confirmed by linear regression modelling using genotype and body mass as independent predictive variables, which revealed a genotype-dependent reduction of lean mass in females not explained by their reduced body mass alone and similar averaged fat amount to controls despite their reduced body mass in both male and females **(Fig. 1e).** Therefore, we conclude that mutant mice are lighter than their littermates mainly because of reduced lean mass, while females have increased fat mass for their size.

**Figure 1.**
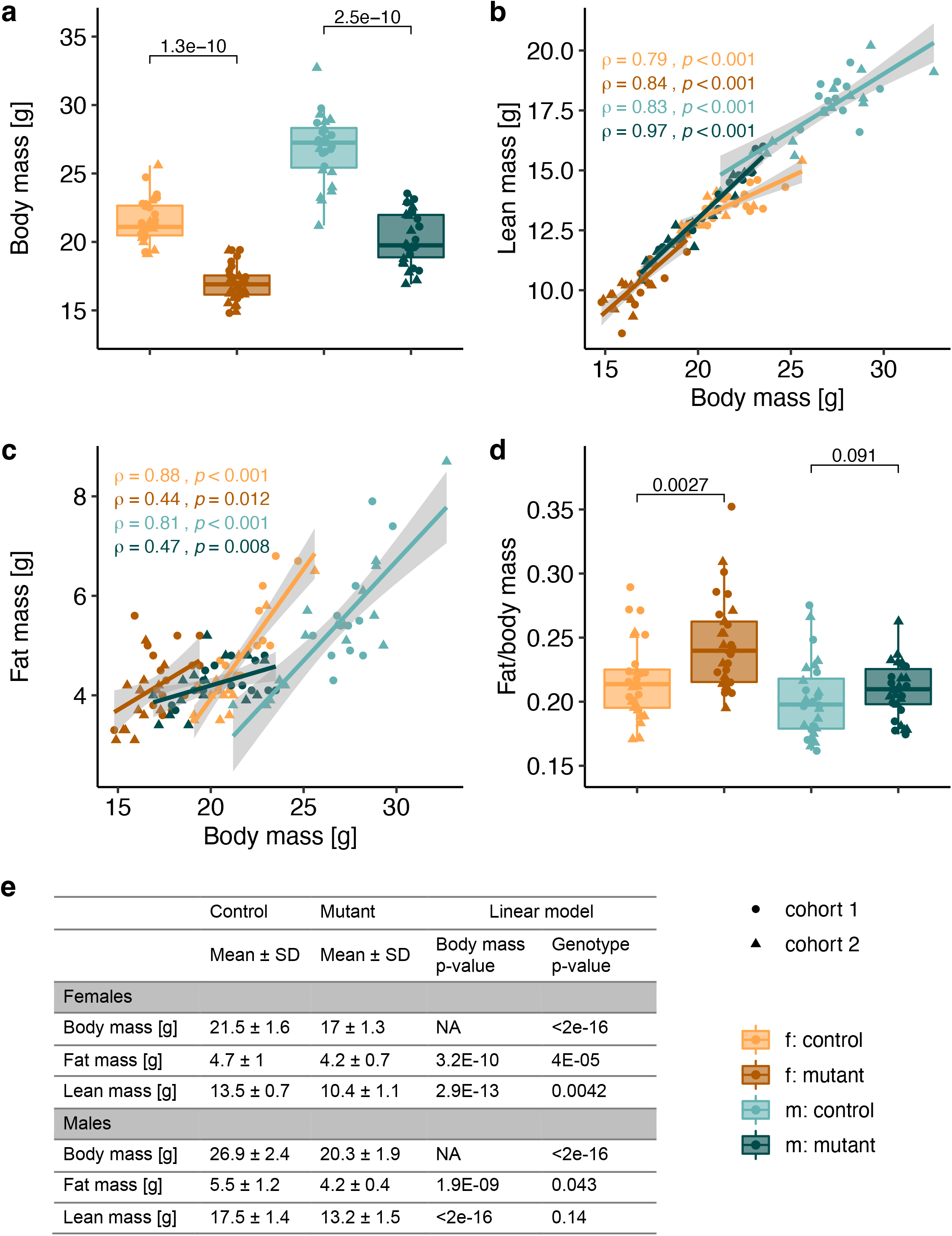
Mutant mice exhibit a shift in body composition. **a** Body weight measured at 14wks old (cohort 1: circles) or 15wks old (cohort 2: triangles). **b-c** Linear regression models: lean mass **(b)** and fat mass **(c),** determined by time domain-nuclear magnetic resonance, in mutant versus control mice using body mass as a covariate. **d** Fat mass of body weight reported as the ratio of fat to relative body weight. **a-d** Each data point represents a single mouse. **e** Summary table of values of body composition parameters. Data are shown as mean ± SD. p-values were determined by linear model. cohort 1: n=14 f. *Mmut*-ki/wt, n=15 f.*Mmut*-ko/ki, n=12 *m.Mmut*-ki/wt, n=15 m.*Mmut*-ko/ki; cohort 2: n=14 f.*Mmut*-ki/wt, n=16 f.*Mmut*-ko/ki, n=16 *m.Mmut*-ki/wt, n=15 m.*Mmut*-ko/ki. (f. females, m. males, mutant: *Mmut*-ko/ki, control *Mmut*-ki/wt)

### Whitening of brown adipose tissue

The observed genotype-dependent increase in fat mass led us to examine the two major fat types. Histological analysis of white adipose tissue (WAT) harvested from the perigonadal region revealed the characteristic unilocular large lipid droplet, containing triglycerides with flattened non-centrally located nucleus in both mutant and control mice. No gross differences between the groups were detected as depicted in **(Fig. 2a).** In contrast, we found clear differences in the appearance of interscapular brown adipose tissue (BAT) between mutant and control mice. Whereas the brown adipocytes of control mice were smaller with a central nucleus and with many (multilocular) lipid droplets, the brown adipocytes of mutant mice showed a unilocular enlarged lipid droplet and displacement of the nucleus to the periphery **(Fig. 2b),** a change observed in all mutants analysed (10/10) but not in the control animals (0/10). This is the so-called BAT whitening process, which has been associated with BAT-dysfunction^12^. This result led us to examine the BAT using an immunohistochemistry marker, the uncoupling protein 1 (UCP1), which is a specific protein able to drive uncoupled respiration for heat production in BAT^15,16^, In fact, we confirmed that despite the morphological changes of brown adipocytes, they showed weak but specific UCP1 protein expression in both sexes **(Fig. 2b).** Consistent with BAT whitening, we found mutant mice to have a significant lower body surface temperature **(Fig. 2c),** suggesting impaired BAT activity results in impaired thermogenesis. We observed significantly reduced plasma leptin concentrations in mutant males compared to controls, with females following the same trend **(Fig. 2d).** As a hormone made predominantly from adipocyte cells, plasma leptin levels strongly correlated with fat mass in linear regression using fat mass as a covariate. Surprisingly, leptin levels were essentially independent of fat mass in *Mmut*-ko/ki mice **(Fig. 2e).** These results suggest leptin production does not correlate with fat amount and it may be dysregulated in mutant animals.

**Figure 2.**
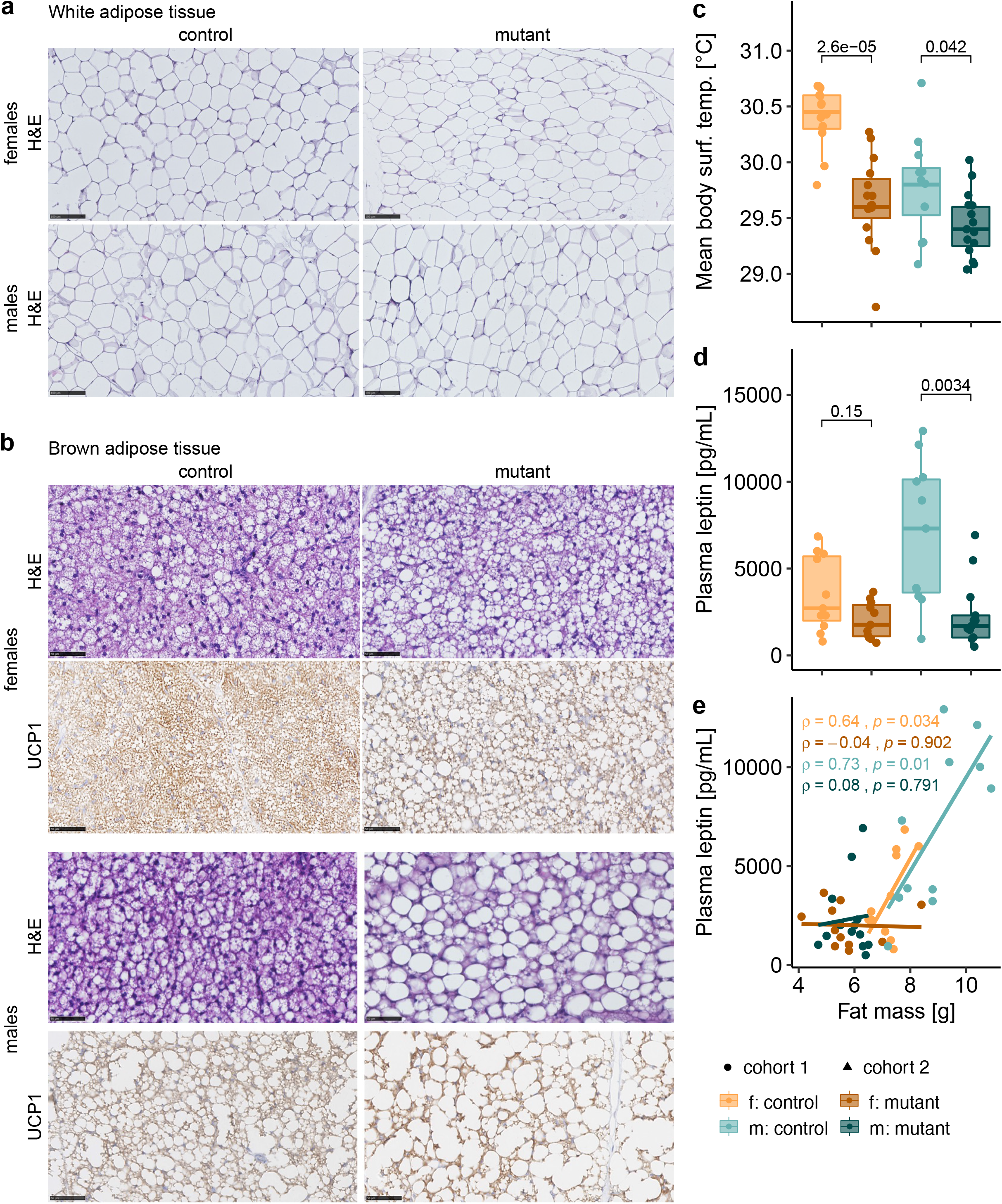
Whitening of brown adipose tissue. **a** Perigonadal white adipose tissue hematoxylin and eosin (H&E) staining of representative control and 4-month-old mutant male and female mice. Scale bar represents 50 μm. **b** Interscapular brown adipose tissue H&E and uncoupling protein 1 (UCP-1) staining of representative 4-month-old control and mutant male and female mice. Scale bar represents 50 μm. **c** Averaged body surface temperature in mice (3-month-old). **d** Plasma leptin levels in mutant mice with littermate controls. **c-d** p-value determined by Wilcoxon test. **e** Plasma leptin levels plotted against fat mass measured through dual-energy X-ray absorptiometry in a linear model (4-month-old). Each data point represents a single mouse. Correlation calculated by the Pearson method. (f. females, m. males, mutant: *Mmut*-ko/ki, control: *Mmut*-ki/wt)

### Reduced adaptation to cold challenge

To further examine how whitening of BAT impacts thermoregulation and energy expenditure, we challenged mice with a short cold exposure. For this experiment, mice spent the first 11 hours under thermoneutral conditions (30 °C) followed by 10 hours at a colder temperature (16 °C). Animals were fasted for the entire trial in order to exclude the influence of food intake; metabolic response was measured by indirect calorimetry. Upon switch from thermoneutral to cold conditions, both mutant and control mice responded by an increased oxygen consumption **(Fig. 3a),** suggesting elevated thermogenesis. Nevertheless, while *Mmut*-ko/ki mice had similar baseline and minimum oxygen consumption under thermoneutral conditions, they were unable to increase their maximum oxygen consumption to the same extent as controls following cold challenge **(Fig. 3a, left panel).** These differences appear to be partly due to their reduced body mass: by linear regression modelling using genotype and body mass as independent predictive variables **(Supplementary Fig. 2a)** we found no difference in baseline oxygen consumption in both females and males, and either no difference (males) or reduced (females) oxygen consumption in mutant mice following cold challenge **(Fig. 3a, right panel).** As expected from their fasted state, both mutant and control mice had a respiratory exchange ratio (RER) that steadily decreased throughout the challenge **(Fig. 3b).** This reduced RER is reflective of a shift towards reduced carbohydrate (CHO) oxidation **(Fig. 3c)** and increased lipid oxidation **(Fig. 3d).** Upon cold challenge, CHO oxidation was not strongly induced in either genotype **(Fig. 3c, left panel),** but female mice did have significantly higher CHO oxidation independent of body mass **(Fig. 3c, right panel, Supplementary Fig. 2b).** By contrast, both mutants and controls had strongly increased lipid oxidation in response to cold challenge **(Fig. 3d, left panel).** Here, mutant mice did not increase lipid oxidation to the same extent as controls **(Fig. 3d left panel);** a difference which is significant in both sexes independent of body mass **(Fig. 3d, right panel, Supplementary Fig. 2c).** Despite their reduced energy production, activity of mutant mice was not significantly reduced compared to controls in both thermoneutral and cold challenge conditions **(Fig. 3e).** Overall, these results reveal that mutant mice show signs of lower metabolism already at thermoneutrality, which is exacerbated by demands on thermoregulation provoked by a cold challenge. Since not all of these differences could be accounted for by adjusting to body mass, other factors (e.g. fat mass, reduced BAT activity) may also contribute to this difference.

**Figure 3.**
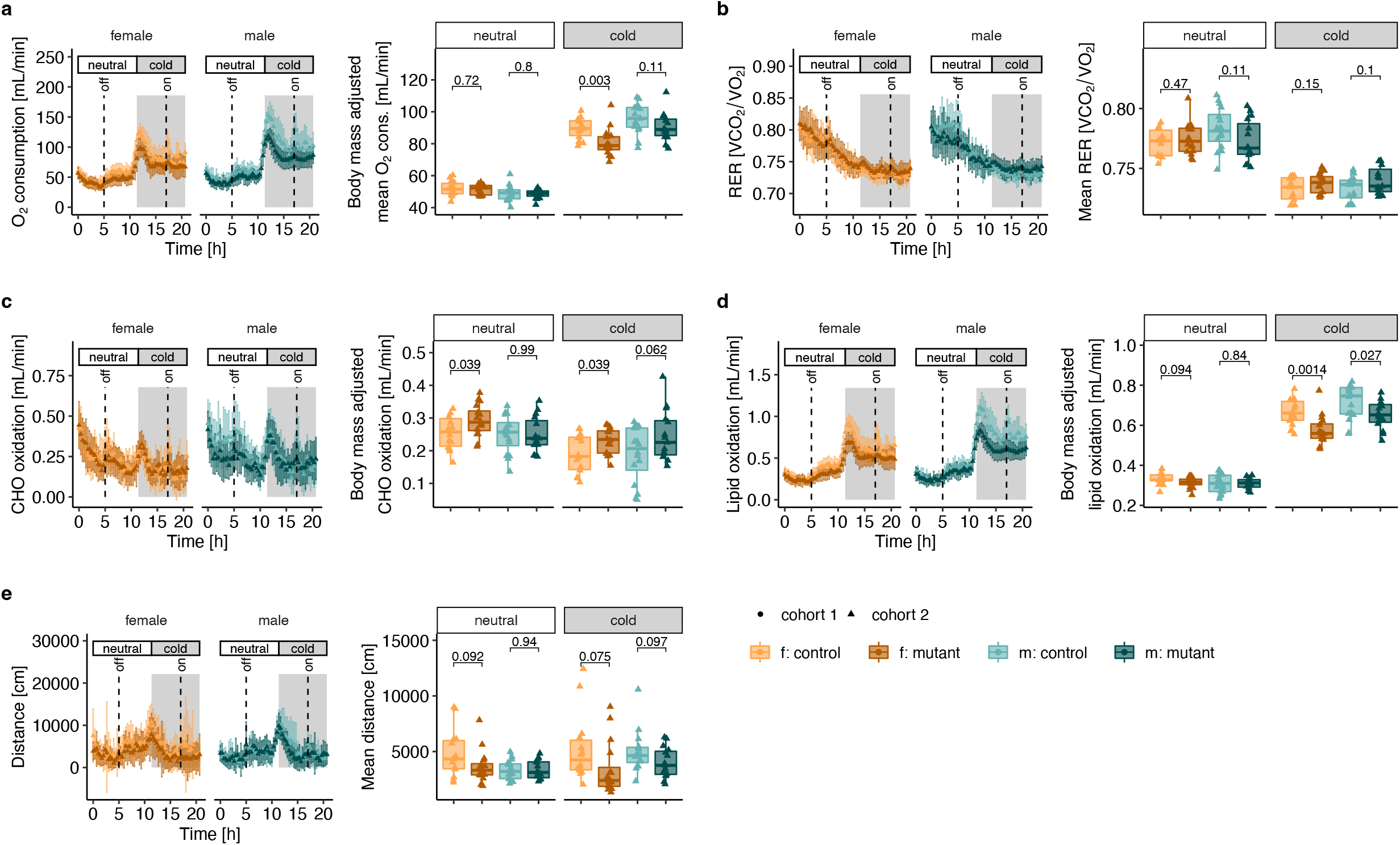
Reduced response to cold challenge. **a** Oxygen consumption, **b** respiratory exchange ratio (RER, ratio of volume of CO_2_ production and O_2_ consumption), c carbohydrate (CHO) oxidation, d lipid oxidation, and e distance covered. Left panels: values obtained following continuous measurement every 20 minutes. Right panels: box plot for each mouse corresponding to residuals from linear correlation with body mass (see **Supplementary Fig. 2).** Mice were housed at thermoneutrality (30°C) from 0 to 10 h and switched to colder temperature (16°C) from 11 to 21 h. Mice were 20-21wks old, fasted, water was provided *ad libitum*. n=14 *Mmut*-ki/wt males and n=16 *Mmut*-ki/wt females; n=16 *Mmut*-ko/ki males and n=15 *Mmut*-ko/ki females, aged 5 months. p-values determined by t-test. (f. females, m. males, mutant: *Mmut-ko/ki*, control: *Mmut*-ki/wt)

### Global hypometabolism

To further study the reduction in metabolism identified during cold challenge, we investigated metabolic function under non-challenging housing conditions (i.e. ambient temperature, *ad libitum* fed). We examined three phases, light phase 1, corresponding to 5 hours before lights off (13:00-18:00, 0 – 5 h), a dark phase following lights off for 12 hours (18:00-06:00, 5 – 17 h), and light phase 2, corresponding to 4 hours following lights on (06:00-10:00, 17 – 21 h) **(Fig. 4).** In this environment, mutant mice showed reduced oxygen consumption, especially at the end of the dark phase and during the light phase 2 **(Fig. 4a).** At the lowest point, the oxygen consumption of mutant females was less than half of their littermates **(Fig. 4a).** The reduced oxygen consumption of mutant mice in the dark phase could not be wholly accounted for by their reduced body mass, as both female and male mutant mice had significantly reduced oxygen consumption when examined using a linear model that included body mass as a predictive variable **(Fig. 4a, Supplementary Fig. 3a).** Periods of reduced oxygen consumption in mutant mice correlated with their lower RER **(Fig. 4b).** This appeared to coincide with an increase in RER in control mice, especially in the dark phase, which was not present in mutant mice **(Fig. 4b).** This RER peak corresponded to enhanced CHO oxidation in control mice in line with the mouse normal feeding behaviour, which was not experienced by mutant mice **(Fig. 4c).** The reduced CHO oxidation in mutant females in the dark phase and light phase 2 was significant independent of their reduced body mass; while body mass appeared to be responsible for the reduced CHO oxidation in mutant males **(Fig. 4c, Supplementary Fig. 3b).** By contrast, lipid oxidation was not changed in mutant males and was slightly increased in mutant females in the dark phase **(Fig. 4d, Supplementary Fig. 3c).** Together, these data suggest the reduced oxygen consumption of females and males in the dark phase are driven by different mechanisms. Despite their reduced energy expenditure, mutant mice appeared similarly active to their control littermates, travelling a similar distance on average and throughout all three measurement periods **(Fig. 4e).** Conversely, mutant mice showed an overall reduced food intake compared to controls **(Fig. 4f),** which started from the “lights off” dark phase coinciding with an expected strong induction of feeding by control animals that was abrogated in mutant mice **(Fig. 4f).** As with CHO oxidation, this difference was independent of body mass only in females **(Fig. 4f, Supplementary Fig. 3d).** Overall, this data suggests that, independent of their decreased body mass, the reduced feeding of females in the dark phase was responsible for their decreased CHO oxidation, which was not replaced by lipid oxidation, culminating in a reduced oxygen consumption. Interestingly, their reduced feeding behaviour was not due to an overall lack of activity. In males, the reduced oxygen consumption in the dark phase were associated with a reduced RER, but could not be ascribed to changes in lipid oxidation, CHO oxidation or food intake when accounting for body mass differences.

**Figure 4.**
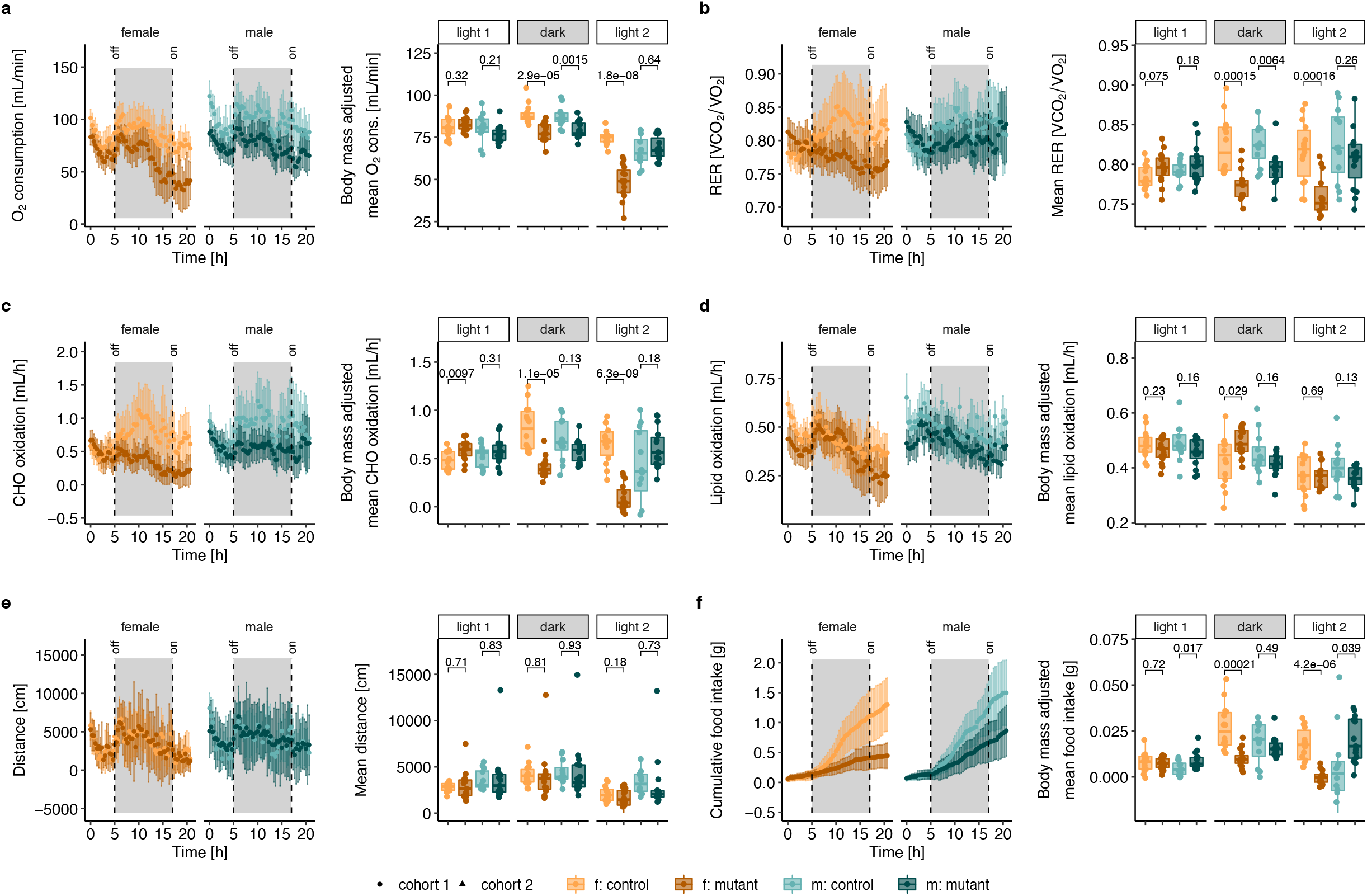
Reduced energy intake and expenditure. **a** Oxygen consumption, **b** respiratory exchange ratio (RER, ratio of volume of CO_2_ production and O_2_ consumption),c carbohydrate (CHO) oxidation, **d** lipid oxidation, **e** distance covered and **f** cumulative food intake (in grams). Left panels: values obtained following continuous measurement every 20 minutes. Right panels: box plot for each mouse corresponding to residuals from linear correlation with body mass (see **Supplementary Fig. 3).** Mice were 14wks old and housed at room temperature (22 °C) with food and water provided *ad libitum* over 21 hours, with data separated according to light phase 1 (0 to 5 h), dark (5 to 17 h), and light phase 2 (17 to 21 h). n=12 *Mmut*-ki/wt males and n=14 *Mmut*-ki/wt females; n=15 *Mmut*-ko/ki males and n=15 *Mmut*-ko/ki females, aged 14 weeks. p-values determined by t-test. (f. females, m. males, mutant: *Mmut-ko/ki*, control: *Mmut*-ki/wt)

### Loss of metabolic flexibility

The reduced ability of male mutant mice to increase their lipid oxidation rate in response to the cold challenge under fasting conditions, along with the poor feeding and reduced carbohydrate oxidation of female mice in standard conditions, suggested a lack of metabolic flexibility and reduced hunger stimulus. This led us to examine plasma concentrations of potential energy sources in both fed and overnight fasted conditions. Compared to controls, mutant female and male mice had reduced glucose concentrations in fed conditions **(Fig. 5a).** However, following fasting, this pattern was reversed. That is, compared to the corresponding control group, fasted mutant mice showed increased glucose concentrations **(Fig. 5a).** These differences appear to be related to an expected reduction in glucose concentrations in response to fasting by control mice, while glucose concentrations in mutant mice stayed approximately constant across both conditions **(Fig. 5a),** indicating constantly low glycemia independent of feeding state. A similar pattern was seen with circulating triglycerides, whereby mutant mice had reduced levels in fed conditions, but similar or even elevated concentrations following fasting **(Fig. 5b).** By contrast, *Mmut*-ko/ki mice displayed decreased cholesterol concentrations compared to controls in both fed and fasted conditions **(Fig. 5c).** The decreased plasma levels of triglycerides and cholesterol of mutant mice in *ad libitum* fed conditions are in line with a constant fatty acid oxidation, despite being fed. Additionally, the elevated glycerol levels of mutant males **(Fig. 5d)** but comparable concentrations of non-esterified fatty acids (NEFA) of both sexes compared to controls **(Fig. 5e)** under fasting conditions also hint towards an increase in lipid utilization, which however was not seen **(Fig. 4d).** Plasma lactate levels were slightly elevated in mutant animals compared to controls, independent of feeding state **(Fig. 5f),** suggestive of glucose metabolization being shifted more towards the anaerobic pathway. Additionally, both control and mutant mice had reduced lactate following fasting compared to *ad libitum* fed condition, potentially reflecting reduced glucose usage in this state^17^. In line with these dysfunctional responses, Fgf21, a key fasting metabolic regulator^18^, was found to be elevated in the plasma of *Mmut*-ko/ki mice in the fed condition **(Fig. 5g).** Although fasting strongly stimulated Fgf21 production in both mutants and controls, such fasting-induced increase was less pronounced in mutant mice **(Fig. 5g).**

**Figure 5.**
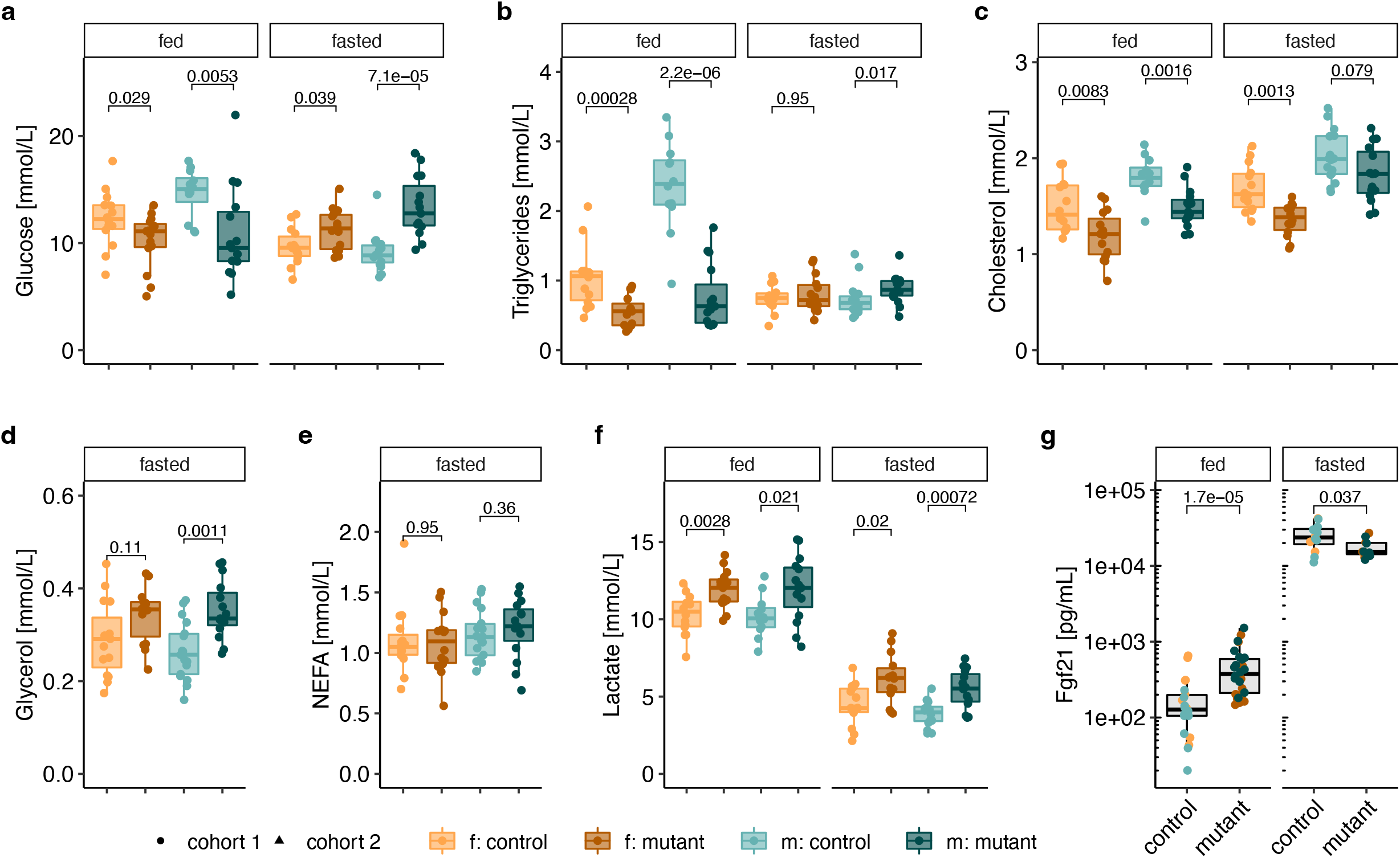
Mutant mice show signs of metabolic inflexibility accompanied by increased levels of Fgf21 when fed. Plasma levels of **a** glucose, **b** triglycerides, **c** cholesterol, **d** glycerol, **e** non-esterified fatty acids (NEFA), **f** lactate and **g** Fgf21 in *ad libitum* fed and/or overnight fasted mice. Plasma levels of Fgf21 were measured from *ad libitum* fed (20wks old) and overnight fasted (18-19wks old) mice. Males and females analyzed together since no sex differences were identified. For all: each data point represents a single mouse, p-values determined by Wilcoxon test. (f. females, m. males, mutant: *Mmut-ko/ki,* control: *Mmut*-ki/wt)

Overall, these changes point toward an incomplete or lack of appropriate response to catabolic conditions, consistent with indirect calorimetry findings. From these data, mutant mice appear to be continuously in a mild, catabolic, “fasted-like” state.

### Impaired glucose homeostasis

The reduction in carbohydrate oxidation under *ad libitum* fed conditions **(Fig. 4),** along with the dysregulated basal plasma glucose and lactate levels in fed and fasted conditions **(Fig. 5),** led us to investigate glucose metabolism after glucose and insulin challenges. We performed intraperitoneal glucose injection as part of a glucose tolerance test in 6 to 7 hour fasted mice **(Fig. 6a).** In these short-term fasted mice, mutant males had reduced glucose concentrations while females were similar to controls **(Fig. 6a).** Following glucose injection, *Mmut*-ko/ki males and females both showed increased circulating glucose levels consisting of elevated peak and residual glucose concentrations across the time course of the experiment (**Fig. 6a).** Combined, these resulted in a significantly higher area under the curve (AUC) for both sexes of mutant mice **(Fig. 6b).** The difference was highest after 30 and 60 minutes, while after two hours mutant animals reached glucose levels only slightly higher than those of controls **(Fig. 6b).** To investigate if the reduced glucose tolerance was a consequence of impaired insulin sensitivity, we performed an intraperitoneal insulin tolerance test, which assesses blood glucose concentration following insulin injection. Prior to this test, again following short (6-7 hours) fasting, *Mmut*-ko/ki mice of both sexes were significantly hypoglycaemic compared to controls **(Supplementary Fig. 4a).** The insulin injection reduced fasting blood glucose levels in *Mmut*-ko/ki mice less efficiently than controls **(Fig. 6c, Supplementary Fig. 4a),** a conclusion supported by higher AUC values in mutant animals **(Fig. 6d),** suggesting a reduced insulin sensitivity in these mice. In this case, the biggest difference in glucose concentrations between control and mutant mice was found in the first 15 minutes following insulin injection, with a strong decrease in glucose levels observed in controls but not mutants at this time point **(Fig. 6d, Supplementary Fig. 4a).** Histological analysis of the pancreas identified similar islet area normalized to pancreatic area between mutants and controls **(Supplementary Fig. 4b),** with no obvious morphological anomalies of the islets **(Supplementary Fig. 4c).** In line with this, immunohistochemistry staining for glucagon and insulin detected by fluorescence **(Supplementary Fig. 4d)** and chromogenic labelling **(Supplementary Fig. 4e)** were consistent with typical size and staining patterns of both hormones in mutants and controls, supportive of normal pancreatic islet function. In conclusion, mutant mice had a reduced glucose tolerance, consistent with their inability to regulate basal glucose levels in fed and fasting conditions. Given the normal islet histology including insulin/glucagon staining as well as the impaired insulin tolerance test, our data points to impaired insulin sensitivity rather than impaired beta cell function as the main cause for the observed glucose intolerance.

**Figure 6.**
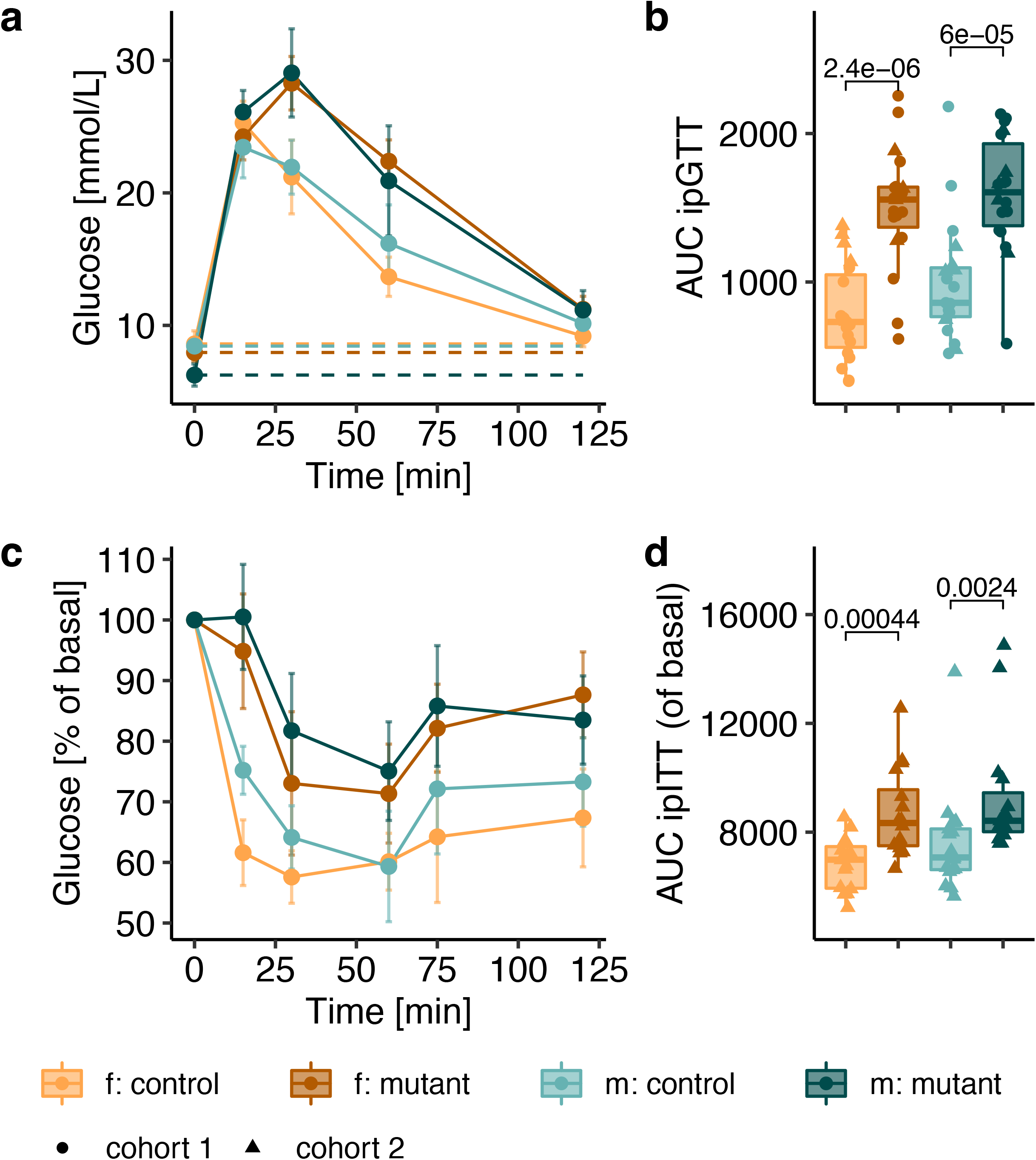
Mutant mice display impaired glucose tolerance. **a** Intraperitoneal glucose tolerance test (ipGTT) curve (data shown as mean ± SD) with baseline glucose shown as a dotted line. **b** Area under the curve (AUC) of glucose excursion shown for time 0-120 minutes. n = 28 *Mmut*-ki/wt females, n=31 *Mmut*-ko/ki females, n=28 *Mmut*-ki/wt males, n=30 *Mmut*-ko/ki males, 14wks old. c. Glucose excursion after intraperitoneal injection of insulin (intraperitoneal insulin tolerance test, ipITT), values are mean ± SD and expressed as percentage of basal (fasting values) (for glucose in [mmol/L] see **Supplementary Fig. 4a). d** Area under the curve of ipITT for time 0-120 minutes. n=15 *Mmut*-ki/wt females, n=16 *Mmut*-ko/ki females, n=17 *Mmut*-ki/wt males, n=16 *Mmut*-ko/ki males, 16wks old. Each data point represents a single mouse. p-values determined by Wilcoxon test. (f. females, m. males, mutant: *Mmut*-ko/ki, control: *Mmut*-ki/wt)

### Liver damage

Since liver contributes to both glucose^19^ and lipid metabolism^20^, we hypothesized that the impaired glucose tolerance observed in *Mmut*-ko/ki animals and the altered lipid profile in plasma may be due to impaired hepatic function, consistent with abnormalities we previously identified in this model^17^. At necropsy, we observed an increased liver weight when normalized to body weight in female *Mut*-ko/ki mice **(Fig. 7a).** This indication of hepatomegaly was associated with signs of liver damage, including slightly elevated plasma levels of alanineaminotransferase (ALAT) and aspartate-aminotransferase (ASAT) in mutant males as well as elevated alkaline phosphatase (ALP) in both sexes **(Fig. 7b).** Relative quantification of methylmalonyl-CoA, the substrate of methylmalonyl-CoA mutase, revealed elevated levels in mutant animals **(Fig. 7c).** Upstream in the metabolic pathway, propionyl-CoA levels were also increased (~15-fold increase), as were those of its derivative metabolite 2-methylcitrate (indistinguishable from methylisocitrate in our assay) (~3-10-fold increase) **(Fig. 7c).** These were consistent with vastly elevated levels of the metabolite methylmalonic acid, quantified in *Mmut*-ko/ki livers **(Fig. 7d),** together suggesting that this damage may have arisen from toxic metabolites^18^. However, electron microscopy did not reveal any ultrastructural differences or signs of mitochondrial pathology in the liver **(Supplementary Fig. 5a).** *Mmut*-ko/ki mouse livers showed normal mitochondria size distribution and ultrastructure, despite variations in mitochondria size and presence of organelles, such as lipid droplets, depending on the sectioning plane. Similar observations were made in respect of control mouse livers. We did not notice any changes in the cristae integrity. Since plasma levels of lactate were strikingly increased in mutant mice **(Fig. 5f),** we hypothesised that the ability of the liver to metabolize lactate into glucose via the gluconeogenesis pathway may be impaired. However, we did not find evidence of impaired gluconeogenesis as determined by an intraperitoneal pyruvate tolerance test in a subset of female mice **(Supplementary Fig. 5b).** Moreover, due to evidence of a constant “fasted-like” state (Fig. 5), we further speculated that these mice may have a defect in glycogen storage and utilization. Nevertheless, we did not identify a qualitative difference in liver glycogen deposition, as determined by histological analysis of periodic-acid Schiff staining in the absence and presence of diastase **(Supplementary Fig. 5c).** Therefore, our results are consistent with significant effects on hepatic metabolism with only mild cellular damage, potentially arising from disease-related metabolites, but do not support reduced gluconeogenesis or dysfunction at the ultrastructural level to be responsible.

**Figure 7.**
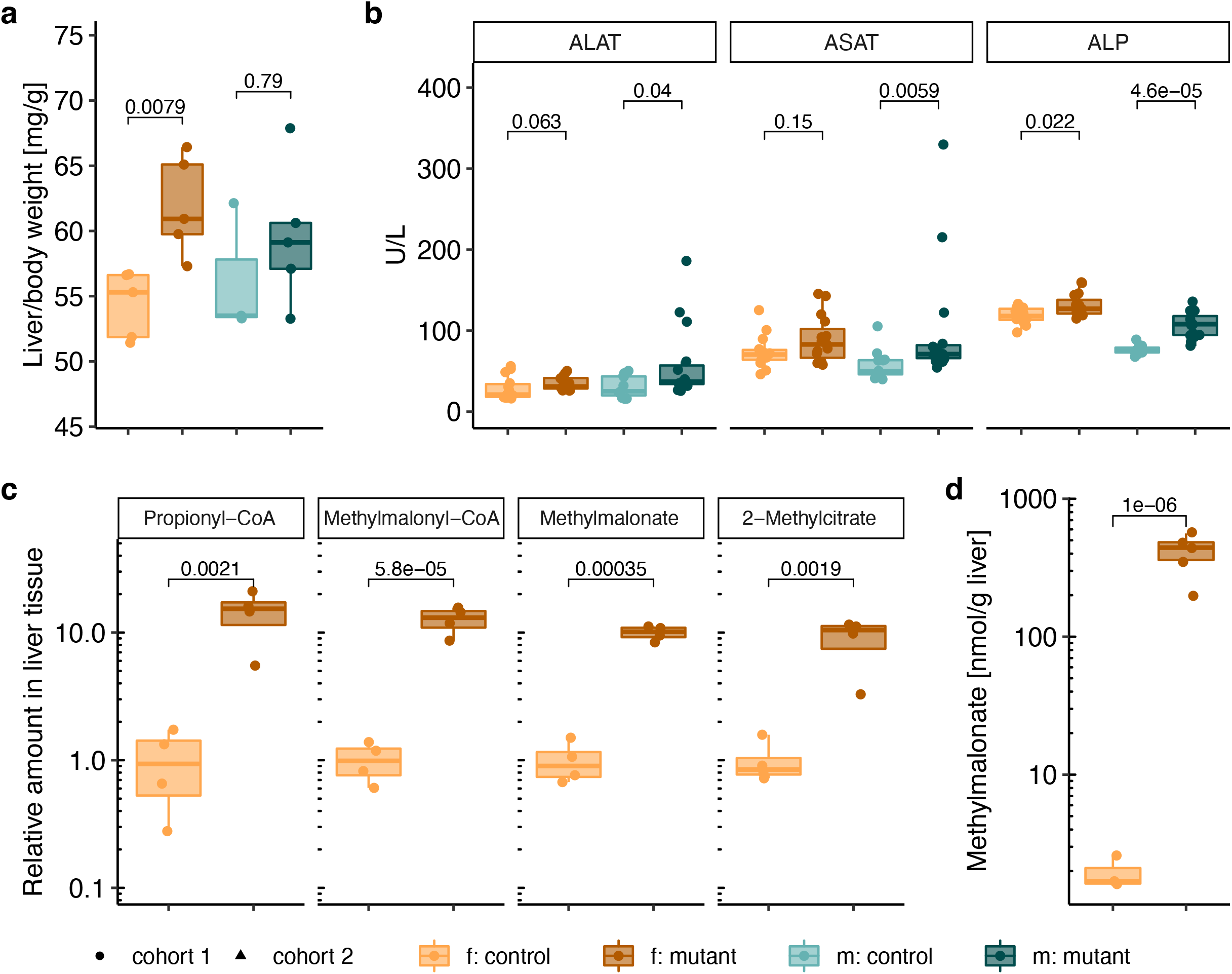
Mutant mice exhibit signs of liver damage. **a** Liver weight normalized to body weight, 20wks old. **b** Plasma levels of alanine-aminotransferase (ALAT/glutamic-pyruvic transaminase (GPT)), aspartate-aminotransferase (ASAT/glutamic-oxaloacetic transaminase (GOT)) and alkaline phosphatase (ALP), 20wks old. **c** Relative levels of propionyl-CoA, methylmalonyl-CoA, methylmalonate and 2-methylcitrate / methylisocitrate in whole liver of females, 6.5-month-old. Data expressed in areas under the curve divided by total ion currents, normalization to the averaged control values. **d** MMA levels in whole liver of females, 6.5-month-old. Each data point represents a single mouse. p-values in a and b determined by Wilcoxon test, in c and d by t-test. (f. females, m. males, mutant: *Mmut*-ko/ki, control: *Mmut*-ki/wt)

### Multi-faceted regulatory changes underlie altered liver metabolism

To investigate the molecular basis of these metabolic changes, we analyzed gene expression by microarray in liver tissue of 9 *Mmut*-ki/wt and 10 *Mmut*-ko/ki male mice in the *ad libitum* fed state. Initial assessment indicated consistent signals across all genes from each sample **(Supplementary Fig. 6a)** and the expected decrease in *Mmut* expression in *Mmut*-ko/ki compared to *Mmut*-ki/wt mice **(Supplementary Fig. 6b),** indicating the data was of good quality. We therefore performed a gene set enrichment analysis, whereby comparison to both Wikipathway^22^ **(Fig. 8a)** and KEGG^23^ **(Supplementary Fig. 6c)** identified positive normalized enrichment score for Ppar signalling pathway-related genes and negative normalized enrichment score for electron-transport chain / oxidative phosphorylation related genes in mutant mice. Mapping of electron transport chain expression changes to individual complexes suggests that this downregulation concerns all complexes of the respiratory chain **(Fig. 8b).** A dysfunction of the respiratory chain would be consistent with the increased blood lactate found previously **(Fig. 5f).** Furthermore, changes in the NAD+/NADH ratio may result in dysregulation of the cellular redox state. Consistent with this, we found altered expression of genes related to glutathione synthesis **(Supplementary Fig. 6d).**

**Figure 8.**
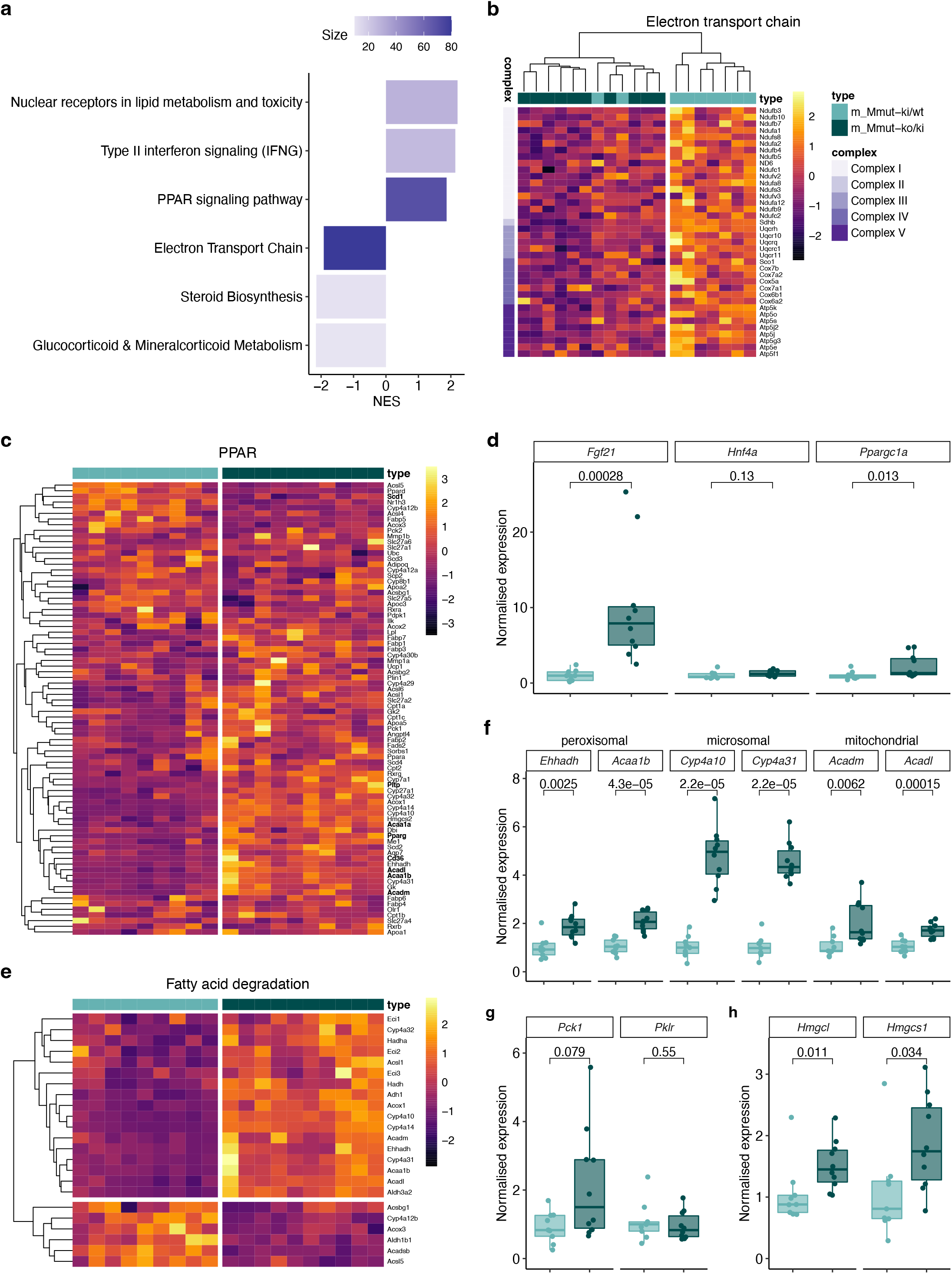
Altered expression of PPAR and electron transport chain-associated genes in mutant mouse liver. **a** WikiPathways gene-set enrichment analysis. **b-c** Heat map of significantly changed genes in electron transport chain **(b)** and PPAR-signaling **(c)** pathways. **d** Confirmatory qRT-PCR of *Fgf21, Hnf4a* and *Ppargc1a* of the same samples. **e** Heat map of significantly changed genes encoding fatty acid degradation proteins. **f-h** Confirmatory qRT-PCR of fatty acid oxidation genes (**f**), rate limiting genes of gluconeogenesis (*Pck1*) and glycolysis (*Pklr*) (**g**), and ketone body synthesis (**h**). All samples derived from RNA from liver tissue of 9 *Mmut*-ki/wt and 10 *Mmut*-ko/ki 4-month-old male mice in the *ad libitum* fed state. All p values determined by Wilcoxon test.

Unsupervised clustering of Ppar-related genes resulted in clear separation of mutant and control samples, and identified Pparγ to be over-expressed in *Mmut*-ko/ki livers **(Fig. 8c).** We observed upregulation of several genes regulated by Pparγ which are associated with metabolism of lipids, including genes involved in lipid transport (e.g. *Pltp*), fatty acid transport (e.g. *Cd36*) and oxidation (e.g. *Bien, Cyp4a1, Acaa1a, Acaa1b, Acadl, Acadm*) as well as lipogenesis (e.g. *Scd-1*) **(Fig. 8c).** Associated to upregulation of Ppar pathways, we confirmed upregulation of *Fgf21* expression in the liver using qRT-PCR **(Fig. 8d),** a finding consistent with increased plasma Fgf21 levels identified in the *ad libitum* state **(Fig. 5g).** *Fgf21* expression can be induced by increased Pparα activity and stimulates fat utilization through the expression of PGC1α (*Ppargc1a*)^19^, whereby PGC1α cooperates with PPARα to stimulate fatty acid oxidation and with HNF4α to stimulate gluconeogenesis^20-22^. Using qRT-PCR, we confirmed increased expression of *Ppargc1a* in mutant mice, as well as no change of *Hnf4a* expression **(Fig. 8d).** Consistent with the role of PGC1α and PPARα to stimulate fatty acid oxidation, we further identified altered expression of genes related to fatty acid oxidation **(Fig. 8e; Supplementary Fig. 6e),** and validated upregulation of known peroxisomal (e.g. *Ehhadh, Acaa1b*), microsomal (e.g. *Cyp4a10, Cyp4a31*) and mitochondrial (e.g. *Acadm, Acadl*) fatty acid oxidation pathway genes by qRT-PCR **(Fig. 8f).** By contrast, consistent with unchanged *Hnf4a* expression **(Fig. 8d),** and no evidence of impaired gluconeogenesis when pyruvate was administered **(Supplementary Fig. 5b),** we found no clear trend to altered expression of genes involved in glucose uptake **(Supplementary Fig. 6f)** or glycolysis and gluconeogenesis **(Supplementary Fig. 6g),** and qRT-PCR confirmed no changes to expression of the main control points of gluconeogenesis (*Pck1*) and glycolysis (*Pklr*) **(Fig. 8g).** Finally, microarray analysis suggested upregulation of genes related to ketone body production **(Supplementary Fig. 6h).** This finding was confirmed by qRT-qPCR, identifying upregulation of *HmgcI* and *Hmgcs1,* two key genes involved in ketone body production **(Fig. 8h).** Interestingly, this may be a reaction to the reduced glycemia in the fed state **(Fig. 5a).**

Overall, these data point to a multi-faceted molecular response in the liver which includes activation of PPAR controlled pathways, including upregulation of *Fgf21* (gene and protein) and *Ppargc1a,* which along with PPARα, shift metabolism towards fatty acid oxidation and away from reliance on glucose in the *ad libitum* fed state.

## Discussion

### Clinical parallels and treatment considerations

In this study, we examined energy balance dysregulation in a mouse model of *mut*-type MMAuria, providing insights into adaptation to chronic energy deficiency. There are also clear parallels between the mouse findings here and those of individuals affected by this disease.

The most obvious phenotype of these mice was their reduced size and weight, identified both here and previously^17,24^. Clinically affected individuals also show consistently poor growth outcomes^7^, a fact for which nutritional factors, particularly over-restriction of natural protein, inadequate protein intake or use of precursor-free amino acids, is often suspected to contribute^25–28^. Since mice in this study were exclusively provided with a high-protein diet, our findings suggest the tendency for smaller stature may not necessarily stem from a reduced protein diet, but rather to be at least partially intrinsic to disease.

We further found a shift in body composition towards less lean and more fat mass amount of body weight, an effect strongest in female mice. Compared to a reference population, female patients with MMAuria have been found to be significantly shorter and to have a higher body mass index, percentage body fat, ratio of abdominal to gluteal circumference, and ratio of central to peripheral body fat^32^. These patients were also found to have decreased resting energy expenditure^33^, similar to the hypometabolism detected in our mouse model. The potential underlying factors for this change towards increased fat accumulation and hypometabolism are manifold, and are discussed further below.

We also found multiple signs of dysregulation in glucose metabolism: mutant mice had hypoglycemia in the *ad libitum* fed and shortly fasted state; hyperglycemia following overnight fasting; and a reduced glucose tolerance that was likely caused by impaired insulin sensitivity. Hypoglycemia is an often cited, if not very well described, complication of MMAuria^31^. On the other hand, diabetic ketosis, a potentially life-threatening metabolic complication characterized by hyperglycemia, ketosis and metabolic acidosis, often associated to type 1 diabetes mellitus^35^, has been described as a rare complication of MMAuria in at least 8 separate case reports^33–40^; with an equal number of patients described in related organic acidurias (reviewed in^36^). In some of these individuals, hyperglycemia was resistant to even large doses of insulin^34,38^. Glucose dysregulation, particularly following bolus administration, is an important consideration in MMAuria disease management, as intravenous glucose is a mainstay of therapy for metabolic decompensation^7^. Our findings suggest that glucose administration must be cautiously dosed under repetitive monitoring of glucose and lactate levels, as the latter may rise when large amounts of glucose are applied^7,41^.

Finally, our findings point to alterations in the liver both at the cellular and molecular levels. Liver changes have been previously reported in MMAuria patients and other mouse models of the disease, including histological abnormalities, mitochondrial pathology, hepatomegaly and more rarely, hepatoblastoma and hepatocellular carcinoma^21,45–50^, suggesting this to be a consistent finding in disease and potentially related to the elevated levels of toxic metabolites. Previous histological investigations conducted on livers of *Mut*-ko/ki mice fed with a 51%-protein diet from day 12 of life revealed morphology changes, such as hepatocellular hypertrophy comprised of enlarged hepatocytes, anisonucleosis, a high number of binucleated forms and a high rate of intranuclear inclusions^13^. Although we did not identify ultrastructural changes here, ultrastructure abnormalities in the mitochondria, such as megamitochondria and disturbed cristae, have been described in patient samples or in other mouse models of MMAuria^45,51–53^. The liver plays an important role in energy balance during fasting and starvation, by regulating carbohydrate and lipid metabolism and by mobilizing energy during nutritional deprivation, it is therefore not surprising that a liver dysfunction would cause energy imbalance in response to fasting^24^. Further, a direct influence of reduced function of MMAuria related enzymes on cholesterol synthesis has recently been described^54^, consistent with our finding of reduced plasma cholesterol in fed and fasted conditions here.

### Potential underlying mechanisms

The pathological features described here represent a global metabolic response to disease pathogenesis and energy challenge, likely mediated through many interconnected pathways. Nevertheless, we identified changes in specific driver molecules which have the capability to enact such a multi-system response and may therefore represent therapeutic targets.

Liver expression and plasma protein levels of Fgf21 were elevated in mutant animals compared to controls in *ad libitum* fed conditions, but fasting-induced increase was less pronounced in mutant mice. Elevated Fgf21 in fed conditions is consistent with findings from another mouse model of MMAuria (Mmut^-/-^;Tg^INS-MCK-*Mmut*^) and MMAuria patients^43,46^ and is positively predictive of muscle mitochondrial pathologies in a quantitative manner^47^. Fgf21 is a metabolic hormone secreted by several organs, including the liver, in response to such factors as starvation, overnutrition and refeeding, low protein, high fat or high carbohydrate diets and methionine restriction^48–53^. Downstream actions controlled by Fgf21 include stimulation of glucose uptake and lipolysis in white adipose tissue, resulting in reduced plasma glucose and circulating triglycerides^48,54^. Therefore, the lower basal plasma glucose and triglyceride levels found in *ad libitum* fed mutant mice may be related to their increased Fgf21. Fgf21 also induces thermogenesis in BAT thereby increasing energy expenditure and improving glucose metabolism^55,56^. In a skeletal muscle *Atg7-ko* mouse, investigators found decreased fat mass along with increased fatty acid oxidation and browning of WAT due to induction of Fgf21^57^. We found quite the opposite: a clear manifestation of BAT “whitening” with worsening glucose tolerance. In our mice, BAT whitening was associated with reduced energy expenditure and body temperature, suggesting impaired thermogenesis despite elevated circulating Fgf21 levels. Our hypothesis is that the process of BAT whitening may be a consequence of an energy-saving mechanism resulting in energy storage in the form of lipids. Thus, we hypothesize that the unilocular enlarged lipid droplets found in BAT and associated reduced body temperature may be due to reduced BAT lipid oxidation. Importantly, BAT whitening is associated with impaired mitochondrial function^58^ and dysfunctional glucose metabolism^68^.

Fgf21 expression is induced by Pparα^49,60^. Pparα, in turn, is a nutritional sensor activated by fatty acid derivates that are formed during lipolysis, lipogenesis or fatty acid catabolism^61^. Ppar signaling, which appears to be activated in fed mutant mice, may have a direct effect on pathways of glucose metabolism – especially on hepatic glucose production^62^. Transcriptomics analysis in liver *of ad libitum* fed animals revealed an upregulation of numerous genes induced by Pparα, suggesting that an increased Pparα activity causes elevated Fgf21 levels. This may be a potential response aimed to upregulate liver fatty acid oxidation in mutant animals in the fed state.

Ppar and Fgf21 further intersect at Ppargc1a, whose hepatic expression is induced by Fgf21. Ppargc1a is a key transcriptional regulator of energy homeostasis, and causes corresponding increases in fatty acid oxidation, tricarboxylic acid cycle flux, and gluconeogenesis without increasing glycogenolysis^19^. In line with this, glucose and triglycerides levels inversely correlated with plasma Fgf21 levels in the fed condition. Thus, it appears as though the reduced lipid oxidation of BAT is matched by an induction in the liver.

## Conclusions

We conclude that our model of MMAuria shows changes which appear poised to combat the effects of chronic energy shortage. These changes are found at every level: whole body (reduced size), tissue (altered fat amount), cell (BAT whitening) and molecular (altered Fgf21/Ppar), with resulting influence on energy expenditure, glucose and fat metabolism. Since many of the findings here have parallels with individuals affected by this disease, as well as related organic acidurias and mitochondriopathies, they have important implications for disease understanding and patient management as well as long-term energetic insufficiency.

## Methods

### Ethics statement

In Zurich, all animal experiments were approved by the legal authorities (license 048/2016; Kantonales Veterinäramt Zürich, Switzerland) and performed according to the legal and ethical requirements. In Munich, all tests performed were approved by the government of Upper Bavaria, Germany (license 046/2016).

### Mouse generation and housing conditions

The experimental *Mut*-ko/ki and the control *Mut*-ki/wt mice were obtained from in-house breedings, by crossing *Mut*-ko/wt females with *Mut*-ki/ki males. These breeders were generated on a C57BL/6N background where mutations were introduced as previously described^24^. Mouse monitoring entailed regular weight measurements. Mice had *ad libitum* access to sterilized drinking water. Starting from day 12 of age, they were fed with a customized diet containing 51% of protein and whose composition was based on the reference diet U8978 version 22 (Safe, France) as previously described^17^. They had *ad libitum* access to this diet, unless otherwise specified under fasting conditions. All mice were bred in Zurich, where they were housed in single-ventilated cages with a 12:12 hour light/dark cycle and an artificial light of approximately 40 Lux in the cage. The animals were kept under controlled humidity (45-55%) and temperature (21 ± 1 °C) and housed in a barrier-protected specific pathogen-free unit. All parameters were monitored continuously. Mice used for the measurements of liver metabolites and enzymes, electron microscopy and pyruvate tolerance test were housed in Zurich until euthanasia, whereas mice used for the other data were shipped to Munich where they had an acclimatization period of 2 weeks. In Munich, mice were housed according to the German Mouse Clinic housing conditions and German laws, and in strict accordance with directive 2010/63/EU. (www.mouseclinic.de). The animals were kept under the same diet in both facilities.

### Metabolite measurements in livers

Mice were anaesthetized with a sedative solution (xylazine 35 mg/kg, ketamine 200 mg/kg, in NaCl 0.9%) administered intraperitoneally, prior to *in vivo* intracardiac perfusion with a washing solution (heparin solution in HBSS buffer, 1%) using a pump. Following harvesting, livers were placed in tissueTUBEs (Covaris) and snap-frozen in liquid nitrogen before cryofracture using the CP02 cryoPREP Automated Dry pulverizer (Covaris) and were then stored in dry ice.

#### Measurement of methylmalonic acid

Liver homogenates were prepared in cold conditions (+4°C) by adding a stainless-steel bead (Qiagen, cat. 69989) and tissue lysis buffer (composition: 250 mM sucrose, 50 mM KCl, 5 mM MgCl2, 20 mM Tris base, pH adjusted to 7.4 with HCl, 5 μL buffer/mg of tissue) to each sample right before mechanical lysis using the TissueLyser II (20 Hz, twice for 90 seconds) (Qiagen). Homogenates were centrifuged at +4°C (600g, 10 min) and supernatants were collected. Protein concentration was assessed (Quick Start Bradford Protein Assay, Bio-Rad) and samples were normalized to ~10 - 25 mg protein/mL. For determination of MMA, 250 - 300 μL of homogenate were used for liquid-liquid extraction. Briefly, 10 μL of 1 mM stable isotopelabelled d3-MMA (Cambridge Isotope Laboratories, Inc., Andover, USA) and 100 μL of 1.25 mM d4-nitrophenol (d4-NP; euriso-top GmbH, Saarbrücken, Germany) were added as internal standards. Samples were acidified with 300 μL of 5 M HCl and after addition of solid sodium chloride extracted twice with 5 mL ethyl acetate each. The combined ethyl acetate fractions were dried over sodium sulfate and then evaporated at 40°C under a stream of nitrogen. Samples were then derivatized with N-methyl-N-(trimethylsilyl)-heptafluorobutyramide (Macherey-Nagel, Düren, Germany) for 1 h at 60°C. For GC/MS analysis, the quadrupole mass spectrometer MSD 5975A (Agilent Technologies, USA) was run in the selective ion-monitoring mode with electron impact ionization. Gas chromatographic separation was achieved on a capillary column (DB-5MS, 30 m x 0.25 mm; film thickness: 0.25 μm; Agilent J&W Scientific, USA) using helium as a carrier gas. A volume of 1μL of the derivatized sample was injected in splitless mode. GC temperature parameters were 80°C for 2 min, ramp 50°C/min to 150°C, then ramp 10°C/min to 300°C. Injector temperature was set to 260°C and interface temperature to 260°C. Fragment ions for quantification were m/z 247 (MMA), m/z 250 (d3-MMA) and m/z 200 (d4-NP). A dwell time of 50 ms was used for d4-NP and 100 ms for MMA and d3-MMA.

#### Measurement of propionyl-CoA, methylmalonyl-CoA, methylmalonate and methylcitrate

500 μL of cold methanol and 350 μL of cold water were added to 25 mg of pulverized frozen tissue in 2 ml tubes containing ceramic beads (1.4 and 2.9 mm). Samples were homogenized using a Precellys evolution tissue homogenizer (Bertin) connected to a cooling system (Cryolysis, Bertin) (3 cycles of 20 sec, 10 000 rpm). A second cycle of homogenization (3 cycles of 20 sec, 10 000 rpm) was performed after addition of 1 mL of 4°C chloroform. Homogenized samples were then centrifuged (16000 x g for 10 min at 4°C). The upper layer (aqueous fraction) was transferred to a new tube and stored at −80°C until LC-MS analysis. Samples were dried down in a vacuum dessicator (Speedvac, Savant) and resuspended in 100 μL of a 50:50 mixture of water and methanol.

LC-MS analysis was performed using a LC-MS qTOF mass spectrometer scanning m/z between 69 and 1700 as described^63^. Briefly, 5 μL of sample was injected and subjected to ion pairing chromatography with an Inertsil 3 μm particle ODS-4 column (150 x 2.1 mm, GL Biosciences) on an Agilent 1290 High-Performance Liquid Chromatography system using hexylamine (Sigma-Aldrich) as pairing-agent. The flow rate was constant at 0.2 mL min-1 using mobile phase A (5 mM hexylamine adjusted with acetic acid to pH 6.3) and B (90% methanol/10% 10 mM ammonium acetate (Biosolve, adjusted to pH 8.5). The buffer gradients and the parameters for detection in an Agilent 6550 ion funnel mass spectrometer operated in negative mode with an electrospray ionization were exactly as described^63^. Metabolite abundance was determined by integrating, in extracted ion chromatograms, peak areas of the predicted m/z with a window of +/-10 ppm. Relative concentrations were determined after normalization to total ion current. Each sample was analysed separately (i.e. the samples were not pooled).

### Assessment of body composition

A body composition analyser (Bruker MiniSpec LF 65) based on time domain-nuclear magnetic resonance (TD-NMR) was used to provide a robust method for the measurement of lean tissue and body fat in live mice without anaesthesia at 13-15 week of age. TD-NMR signals from all protons in the entire sample volume were used and data on lean and fat mass was provided. Body composition analysis was based on a linear regression modelling using body mass or lean mass as a covariate, as previously recommended^73^. Fat mass assessed using the dual energy X-ray absorption analyser UltraFocus^DXA^ by Faxitron®, in anaesthetised mice, at the age of 20 weeks was used for correlation plot with serum leptin levels.

### Body surface temperature

Body surface temperature was measured using Infrared thermography, which is a passive, remote, and non-invasive method. We use a high-resolution A655sc FLIR infrared camera and process the data with FLIR ResearIR Max software. A mouse is isolated, and least three images per mouse are taken when eyes visible and their four feet are on the floor. The average of the whole-body surface temperature and the maximum temperature (eye) is used to evaluate the phenotype^65–68^. Alopecia, sex and age^78^, stress parameter/behaviour^66,70^ and metabolism^80^, among others, represent some of the factors affecting the surface temperature of the mouse. Since skin problems and over-grooming behaviour are the most important reasons for hair loss, which is observed very frequently in C57BL substrains^72^, finally influencing the outcomes, we have scored the hair loss in order to control confounding effects.

### Indirect calorimetry

Indirect calorimetry was performed as described^73^. Briefly, high precision CO_2_ and O_2_ sensors measured the difference in CO_2_ and O_2_ concentrations in air flowing through control and animal cages. The rate of oxygen consumption took into account the air flow through the cages that was measured in parallel. Data for oxygen consumption were expressed as ml O_2_*h^−1^*animal^-1^. The system also monitored CO_2_ production; therefore, the respiratory exchange ratio (RER) could be calculated as the ratio VCO_2_/VO_2_. The test was performed at room temperature (23°C) with a 12:12 hours light/dark cycle in the room (lights on 06:00 CET, lights off 18:00 CET). Wood shavings and paper tissue were provided as bedding material. Each mouse was placed individually in a chamber for a period of 21 hours (from 13:00 CET to 10:00 CET next day) with free access to food and water. Metabolic cuvettes were set up in a ventilated climate room continuously supplied with fresh air from outside. The activity was measured using light beam frames on an x and y axes. The parameter named Distance was calculated from x and y counts including ambulatory activity, fine movements and total activity. The carbo and lipid oxidation rates were calculated using the Frayn formulas: CHO oxidation [g/min] (4.55*VCO2-3.21*VO2), Lipid oxidation [g/min]: (1.67*VO2-1.67*VCO2) with the protein oxidation considered negligible. VO_2_ and VCO_2_ were converted in grams.

For VO2, CHO and lipid oxidation we used the body weight-independent residuals since body mass is the major determinant for variability in absolute VO2, CO2, CHO and lipid oxidation, food intake. Therefore, we calculated a linear model as previously described^82,83^. Briefly, the residuals of these models represent the differences between the observed individual values and predicted ones by their mean body weight (body weight before and after the indirect calorimetry trial).

### Collection of blood samples

Blood samples were collected by puncture of the retrobulbar plexus under isoflurane anesthesia in Li-heparin coated sample tubes from two cohorts of mice: In one cohort samples from overnight-fasted mice were collected at 12 weeks of age and samples of *ad libitum* fed mice at the age of 20 weeks. In the second cohort samples from *ad libitum* fed animals were collected at 9-11 weeks of age and samples after overnight fasting at 18-19 weeks. Blood samples collected after overnight food withdrawal were cooled in a rack on ice after collection.., while samples collected from fed animals were stored at room temperature. Samples were separated within one hour or after 1-2 hours by centrifugation (5000xg, 10min, 8°C) for fasting and fed condition, respectively, and plasma was transferred to 1.5ml conic sample tubes and immediately analysed for clinical chemistry. Aliquots for biomarker measurements Samples collected from *ad libitum* fed mice were stored at room temperature until separation within 1-2 hours by centrifugation and transfer of plasma to 1.5ml sample tubes for clinical analysis and were transferred to 96-well micro-well plates. for biomarker measurements.

### Plasma clinical chemistry and biomarker analysis

Plasma lipid and glucose levels were determined using an AU480 clinical chemistry analyser (Beckman-Coulter) and adapted reagents from Beckman-Coulter (glucose, cholesterol, triglycerides and lactate) evtl Wako-Chemicals (NEFA) and Randox (Glycerol) as previously described^73^. We use a multiplex assay platform to measure the concentration of leptin, FGF21 and Insulin in plasma samples. It is an electroluminescence-linked immunosorbent assay based on the Mesoscale Discovery technology (U-Plex, Mesoscale Diagnostics, Rockville, Maryland, USA). A 10 spot MSD plate is coated with anti-insulin, anti-FGF21 and anti-leptin antibodies (previously treated with the corresponding spot-linkers). Plasma samples are diluted 1:2 and incubated for 1h. After that, the samples are incubated for 1 hour with Sulfotag conjugated detection antibodies (second antibodies) before they are analysed in the MSD plate reader. Coated and detection antibodies for leptin, FGF21 and insulin are provided as U-plex antibodies from MSD. MSD discovery workbench is used as analysis software.

### Intraperitoneal glucose tolerance test

Glucose metabolism disturbance was determined using the glucose tolerance test (GTT). Glucose was administered intraperitoneally (i.p.) after a 6-7h food withdrawal and glucose levels were measured at specific time points in the subsequent 2 hours using blood drops collected from the tail vein. The body weight of mice was determined before and after food withdrawal. Basal fasting blood glucose level was analyzed with the Accu-Chek Aviva Connect glucose analyzer (Roche/Mannheim). Thereafter, glucose was injected i.p. (2 g/kg) and circulating glucose was measured 15, 30, 60, and 120 minutes later.

### Insulin tolerance test

The insulin tolerance test was performed in mice after a 6-7 hours-lasting food withdrawal. At the beginning of the test, the body weight of mice was measured.Body weight was measured and fasting blood glucose levels were assessed from a drop of blood collected from the tail vein using a hand-held glucometer (AccuCheck Aviva, Roche, Mannheim, Germany). Insulin (0.75 U/kg body weight Huminsulin® Normal 100, Lilly) was injected intraperitoneally using a 25-gauge needle and a 1-mL syringe and after 15, 30, 60, 75 and 120 minutes additional blood samples were collected and used to determine blood glucose levels as described before. Repeated bleeding was induced by removing the clot from the first incision and slightly massaging the mouse tail. Mice were not given any food during the test but after the experiment was finished, they were placed in a cage with plentiful supply of water and food.

### Pyruvate tolerance test

The pyruvate tolerance test was performed in mice after a 6-7 hours-lasting food withdrawal. At the beginning of the test, the body weight of mice was measured. Blood was collected from tail vein without restraining the animal and using a scalpel blade. Fasting blood glucose levels were assessed using the Accu-Chek Aviva meter and test strips (Roche) in 0.6 μL of blood. Thereafter mice were injected intraperitoneally with 2 g of sodium pyruvate (Sigma)/kg body weight) using a 20 % sodium pyruvate solution (in 0.9% NaCl), a 25-gauge needle and a 1-mL syringe. 15, 30, 45, 60, 90 and 120 minutes after sodium pyruvate injection, additional blood samples (two drops each to have duplicates) were collected and used to determine blood glucose levels as described before. Repeated bleeding was induced by removing the clot from the first incision and slightly massaging the mouse tail. Mice were not given any food during the test but after the experiment was finished, they were placed in a cage with plentiful supply of water and food. 8 *Mut*-ki/wt and 8 *Mut*-ko/ki females were used initially, but two animals (one per group) were identified as outliers and removed from the dataset.

### Electron microscopy

Mice were anaesthetized with a sedative solution (xylazine 35 mg/kg, ketamine 200 mg/kg, in NaCl 0.9%) administered intraperitoneally, prior to *in vivo* intracardiac perfusion using a pump. Following a perfusion with 1 % heparin in Hanks’ Balanced Salt solution to wash out the blood, mice were perfused with a Working Solution (2 % formaldehyde (powder Sigma #158127), 0.01% glutaraldehyde (ampoules EMS #16220 EM Grade), in 0.1 M caco-sucrose buffer (0.2 M pH 7.4 with cacodylate C_2_H_6_AsNaO_2_*3H_2_O, powder Merck #8.20670.0250), in double-distilled water). The Working Solution was filtered with a 0.22 μm filter prior to use. Following harvesting, livers were washed twice in Working Solution and stored in fixative storage solution (0.1 M Cacodylate-sucrose buffer, 2.5 % glutaraldehyde, in ddH_2_O) before sectioning with a vibratome (Vibratome Series 1000, Ted Pella Inc, Redding CA, USA). Vibratome sections or small dissected pieces (ca. 1 mm3) were subsequently rinsed with 0.1 M cacodylate buffer (pH 7.4) three times for 5 minutes, post-fixed with 1% OsO4 for 1 hour in 0.1 M cacodylate buffer at 0°C, rinsed with H_2_O three times for 5 minutes, block stained with 1 % aqueous uranylacetate for 1 hour, and rinsed with H_2_O twice for 5 minutes. Samples were then dehydrated in an ethanol series and embedded in Epon/Araldite (Sigma-Aldrich, Buchs, Switzerland). Ultrathin (70 nm) sections were post-stained with Reynolds lead citrate and examined with a CM100 transmission electron microscope (Thermo Fisher Scientific, Eindhoven, The Netherlands) at an acceleration voltage of 80 kV using an Orius 1000 digital camera (Gatan, Munich, Germany), or a Talos 120 transmission electron microscope at an acceleration voltage of 120 kV using a Ceta digital camera and the MAPS software package (Thermo Fisher Scientific, Eindhoven, The Netherlands). Pictures were analysed using ImageJ 1. 52p.

### Histopathological analysis

We analysed a total of 40 animals, 20 females (10 *Mmut*-ko/ki and 10 *Mmut*-ki/wt) and 20 males (10 *Mmut*-ko/ki and 10 *Mmut*-ki/wt) in two cohorts at the age of 20 weeks. They were euthanized with CO_2_ and the visceral organs were analysed macroscopically and weighed. For microscopic analyses, we used 4% formalin-fixed buffered paraffin-embedded (3 μm) sections stained with hematoxylin and eosin (H&E).

In pancreas, liver and fat tissues special stains were performed. For the liver, we used serial sections stained with the PAS (periodic acid-Schiff) reaction. This staining method is used to detect polysaccharides such as glycogen, which normally give a magenta color (called PAS positive). To confirm that the staining is due to the presence of glycogen and no other polysaccharides, we compared a section of PAS digested with diastase (which must be PAS negative) with its serial section of PAS (which must be positive).

Using H&E, we examined the two main types of fat, white adipose tissue (WAT) and brown adipose tissue (BAT), collected from the perigonadal and interscapular regions, respectively. In the pancreas we quantified the area of the islet cells normalized by the total area of the pancreatic section.

Immunohistochemical staining in pancreas and fat sections was carried out on 1-2 μm section cut from paraffin blocks in an automated immunostainer from Leica biosystems (BOND RX, using the ds9800-DAB Polymer Detection System), a kit for the streptavidin-peroxidase method. If not otherwise mentioned heat induced antigen retrieval with citrate buffer (pH 6) was performed prior to incubation with the following rabbit mono or poly-clonal antibodies: UCP1 (Abcam ab10983) 1:800, Insulin (cell signaling technology, Catalog Nr. 3014) 1:12.000, and Glucagon (Sigma, Catalog. Nr. K79BB10) 1:2000. For the Immunofluorescence (IF) we used same anti-insulin and anti-glucagon primary antibodies but as secondary antibody we used Goat anti-Rabbit IgG (H+L) Cross-Adsorbed Alexa Fluor 488 (ThermoFischer scientific, Catalog Nr. A-11008) and Alexa Fluor 647 (ThermoFischer scientific, Catalog Nr. A-21244) both in a concentration of 4μg/mL. To confirm antibody specificity positive controls with known protein expression as well as negative controls without primary antibody were used. The slides were analysed by two pathologists independently.

### Molecular phenotyping

Genome-wide transcriptome analysis was performed on ten mutant and nine control male mice. Briefly, total RNA was isolated employing the RNeasy Mini kit (Qiagen) including Trizol treatment and RNA quality assessed by an Agilent 2100 Bioanalyzer (Agilent RNA 6000 Pico Kit). RNA was amplified using the WT PLUS Reagent Kit (Thermo Fisher Scientific Inc., Waltham, USA). Amplified cDNA was hybridized on Mouse Clariom S arrays (Thermo Fisher Scientific). Staining and scanning (GeneChip Scanner 3000 7G) was performed according to manufacturer’s instructions. Transcriptome Analysis Console (TAC; version 4.0.0.25; Thermo Fisher Scientific) was used for quality control and to obtain annotated normalized SST-RMA gene-level data.

Real-time qRT–PCR was performed on cDNA amplified by PrimeScript II 1st Strand cDNA Synthesis Kit (Takara) using GoTaq qPCR (Promega) following manufacturer’s protocols. The products were analyzed on a LightCycler 480 System (Roche). Mouse *Actb* was used as a housekeeping gene for normalization, and relative target gene expression was quantified by the 2^-ΔΔCt^ method. Primer sequences used are listed in **Supplementary Table 1.**

### Statistical analysis and data availability

Data analysis was performed using R (version 4.1.0). Tests for genotype effects were made by using t-test and Wilcoxon rank sum tests. Certain parameters (see figure legends) have been corrected for body weight effects using linear models. A p-value <0.05 has been used as level of significance. All raw data and R scripts are available through an online repository at https://github.com/pforny/MetabolicSwitchMMA. Array data has been submitted to the GEO database at NCBI (GSE188931).

## Acknowledgements

This work received financial support from the University Research Priority Program of the University of Zurich (URPP) ITINERARE—Innovative Therapies in Rare Diseases, and was supported by the Rare Disease Initiative Zurich (radiz), a clinical research priority program for rare diseases of the University of Zurich. M.R.B. is supported by the Swiss National Science Foundation [31003A_175779] and the Wolferman Nägeli Foundation; D.S.F. is supported by the Swiss National Science Foundation [310030_192505]; MHdA is supported by the German Federal Ministry of Education and Research [Infrafrontier grant 01KX1012] and the German Center for Diabetes Research (DZD e.V.). P.F. is supported by the Filling the Gap grant, Medical Faculty, University of Zurich, Switzerland. The authors declare that they have no conflicts of interest with the contents of this article. We thank Martin Poms and Ivan Hartling (Division of Clinical Chemistry and Biochemistry, University Children’s Hospital Zurich) for assistance in data analysis, José María Mateos (Center for Microscopy and Image Analysis, University of Zurich) for his advice about electron microscopy, and GMC technicians and animal caretakers for their expert technical support.

## Author contributions

**M.L.** contributed to the design of the study, generation and monitoring of the animals, scientific coordination, generation, analysis and interpretation of data, and writing the manuscript.

**R.G.** contributed to analysis and interpretation of data (Figures 3 and 4, Suppl. Tables 1 and 2) and writing the manuscript.

**B.R.** contributed to planning the experiments, generation, analysis and interpretation of data (Figure 5) and writing the manuscript.

**J.C-W.** contributed to analysis and interpretation of data (Figure 2; Suppl. Figures 1, 2 and 3) and writing the manuscript.

**P.F.** contributed to interpretation of data, creation of the figures, and reviewing the manuscript.

**S. W.** contributed to technical and scientific input on the whole dataset, and reviewing the manuscript.

**A. K.** contributed to data generation and interpretation (Suppl. Figure 3.)

**F. T.** contributed to data generation and interpretation (Figure 8).

**M. F.** and **C. B.** contributed to mouse genotyping.

**A. A-P.** contributed to data generation and interpretation (Figure 2 and 5).

**M.I.** contributed to generation and interpretation of Figure 8.

**J.B.** contributed to discussion of data and contributed funding

**S. S.** and **S. K.** contributed to data generation and interpretation (Figure 7).

**J.P.D.** contributed to planning the experiments, data generation and interpretation (Figure 7).

**G.T.B.** contributed to planning the experiments (Figure 7), and reviewing the manuscript.

**D.H.** contributed to technical and scientific input (Figures 3 and 4)

**V.G-D.** contributed in conceptualising and supervision of the mouse phenotyping **H. F.** contributed in conceptualising and supervision of the mouse phenotyping

**J.R.** contributed to planning the experiments, data generation and analysis (Figure 3).

**D.S.F.** conceived the idea of the project, contributed to the interpretation of the whole dataset and to the writing of the manuscript.

**M.R.B** conceived the idea of the project

**M.R.H.A.** conceived the mouse phenotyping, funding acquisition

## References

1. Smith, R.L., Soeters, M.R., Wust, R.C.I. & Houtkooper, R.H. Metabolic Flexibility as an Adaptation to Energy Resources and Requirements in Health and Disease. Endocr Rev 39, 489–517 (2018).

2. Kolker, S. et al. Methylmalonic acid, a biochemical hallmark of methylmalonic acidurias but no inhibitor of mitochondrial respiratory chain. J Biol Chem 278, 47388–93 (2003).

3. Luciani, A., Denley, M.C.S., Govers, L.P., Sorrentino, V. & Froese, D.S. Mitochondrial disease, mitophagy, and cellular distress in methylmalonic acidemia. Cell Mol Life Sci 78, 6851–6867 (2021).

4. Schwab, M.A. et al. Secondary mitochondrial dysfunction in propionic aciduria: a pathogenic role for endogenous mitochondrial toxins. Biochem J 398, 107–12 (2006).

5. Okun, J.G. et al. Neurodegeneration in methylmalonic aciduria involves inhibition of complex II and the tricarboxylic acid cycle, and synergistically acting excitotoxicity. J Biol Chem 277, 14674–80 (2002).

6. Coude, F.X., Sweetman, L. & Nyhan, W.L. Inhibition by propionyl-coenzyme A of N-acetylglutamate synthetase in rat liver mitochondria. A possible explanation for hyperammonemia in propionic and methylmalonic acidemia. J Clin Invest 64, 1544–51 (1979).

7. Forny, P. et al. Guidelines for the diagnosis and management of methylmalonic acidaemia and propionic acidaemia: First revision. J Inherit Metab Dis (2021).

8. Kolker, S. et al. The phenotypic spectrum of organic acidurias and urea cycle disorders. Part 1: the initial presentation. J Inherit Metab Dis 38, 1041–57 (2015).

9. Horster, F. et al. Long-term outcome in methylmalonic acidurias is influenced by the underlying defect (mut0, mut-, cblA, cblB). Pediatr Res 62, 225–30 (2007).

10. Peters, H. et al. A knock-out mouse model for methylmalonic aciduria resulting in neonatal lethality. J Biol Chem 278, 52909–13 (2003).

11. Chandler, R.J. et al. Metabolic phenotype of methylmalonic acidemia in mice and humans: the role of skeletal muscle. BMC Med Genet 8, 64 (2007).

12. Shimizu, I. & Walsh, K. The Whitening of Brown Fat and Its Implications for Weight Management in Obesity. Curr Obes Rep 4, 224–9 (2015).

13. Freminet, A., Leclerc, L., Gentil, M. & Poyart, C. Effect of fasting on the rates of lactate turnover and oxidation in rats. FEBS Lett 60, 431–4 (1975).

14. Fisher, F.M. & Maratos-Flier, E. Understanding the Physiology of FGF21. Annu Rev Physiol 78, 223–41 (2016).

15. Petersen, M.C., Vatner, D.F. & Shulman, G.I. Regulation of hepatic glucose metabolism in health and disease. Nat Rev Endocrinol 13, 572–587 (2017).

16. Nguyen, P. et al. Liver lipid metabolism. J Anim Physiol Anim Nutr (Berl) 92, 272–83 (2008).

17. Lucienne, M. et al. In-depth phenotyping reveals common and novel disease symptoms in a hemizygous knock-in mouse model (Mut-ko/ki) of mut-type methylmalonic aciduria. Biochim Biophys Acta Mol Basis Dis 1866, 165622 (2020).

18. de Keyzer, Y. et al. Multiple OXPHOS deficiency in the liver, kidney, heart, and skeletal muscle of patients with methylmalonic aciduria and propionic aciduria. Pediatr Res 66, 91–5 (2009).

19. Potthoff, M.J. et al. FGF21 induces PGC-1alpha and regulates carbohydrate and fatty acid metabolism during the adaptive starvation response. Proc Natl Acad Sci U S A 106, 10853–8 (2009).

20. Lustig, Y. et al. Separation of the gluconeogenic and mitochondrial functions of PGC-l{alpha} through S6 kinase. Genes Dev 25, 1232–44 (2011).

21. Koo, S.H. et al. PGC-1 promotes insulin resistance in liver through PPAR-alpha-dependent induction of TRB-3. Nat Med 10, 530–4 (2004).

22. Rhee, J. et al. Regulation of hepatic fasting response by PPARgamma coactivator-1alpha (PGC-1): requirement for hepatocyte nuclear factor 4alpha in gluconeogenesis. Proc Natl Acad Sci U S A 100, 4012–7 (2003).

23. Martens, M. et al. WikiPathways: connecting communities. Nucleic Acids Res 49, D613–D621 (2021).

24. Forny, P. et al. Novel Mouse Models of Methylmalonic Aciduria Recapitulate Phenotypic Traits with a Genetic Dosage Effect. J Biol Chem 291, 20563–73 (2016).

25. Touati, G. et al. Methylmalonic and propionic acidurias: management without or with a few supplements of specific amino acid mixture. J Inherit Metab Dis 29, 288–98 (2006).

26. Manoli, I., Myles, J.G., Sloan, J.L., Shchelochkov, O.A. & Venditti, C.P. A critical reappraisal of dietary practices in methylmalonic acidemia raises concerns about the safety of medical foods. Part 1: isolated methylmalonic acidemias. Genet Med 18, 386–95 (2016).

27. Evans, M., Truby, H. & Boneh, A. The Relationship between Dietary Intake, Growth, and Body Composition in Inborn Errors of Intermediary Protein Metabolism. J Pediatr 188, 163–172 (2017).

28. van der Meer, S.B. et al. Clinical outcome of long-term management of patients with vitamin B12-unresponsive methylmalonic acidemia. J Pediatr 125, 903–8 (1994).

29. Hauser, N.S., Manoli, I., Graf, J.C., Sloan, J. & Venditti, C.P. Variable dietary management of methylmalonic acidemia: metabolic and energetic correlations. Am J Clin Nutr 93, 47–56 (2011).

30. Feillet, F., Bodamer, O.A., Dixon, M.A., Sequeira, S. & Leonard, J.V. Resting energy expenditure in disorders of propionate metabolism. J Pediatr 136, 659–63 (2000).

31. Halperin, M.L., Schiller, C.M. & Fritz, I.B. The inhibition by methylmalonic acid of malate transport by the dicarboxylate carrier in rat liver mitochondria. A possible explantation for hypoglycemia in methylmalonic aciduria. J Clin Invest 50, 2276–82 (1971).

32. Rugg-Gunn, C.E., Deakin, M. & Hawcutt, D.B. Update and harmonisation of guidance for the management of diabetic ketoacidosis in children and young people in the UK. BMJ Paediatr Open 5, e001079 (2021).

33. Dejkhamron, P. et al. Isolated methylmalonic acidemia with unusual presentation mimicking diabetic ketoacidosis. J Pediatr Endocrinol Metab 29, 373–8 (2016).

34. Boeckx, R.L. & Hicks, J.M. Methylmalonic acidemia with the unusual complication of severe hyperglycemia. Clin Chem 28, 1801–3 (1982).

35. Guven, A. et al. Methylmalonic acidemia mimicking diabetic ketoacidosis in an infant. Pediatr Diabetes 13, e22–5 (2012).

36. Sharda, S., Angurana, S.K., Walia, M. & Attri, S. Defect of cobalamin intracellular metabolism presenting as diabetic ketoacidosis: a rare manifestation. JIMD Rep 11, 43–7 (2013).

37. Mathew, P.M. & Hamdan, J.A. Transient diabetes mellitus in neonatal methylmalonic aciduria. J Inherit Metab Dis 11, 218–9 (1988).

38. Filippi, L. et al. Insulin-resistant hyperglycaemia complicating neonatal onset of methylmalonic and propionic acidaemias. J Inherit Metab Dis 32 Suppl 1, S179–86 (2009).

39. Ciani, F. et al. Lethal late onset cblB methylmalonic aciduria. Crit Care Med 28, 2119–21 (2000).

40. Imen, M. et al. Methylmalonic acidemia and hyperglycemia: an unusual association. Brain Dev 34, 113–4 (2012).

41. Baumgartner, M.R. et al. Proposed guidelines for the diagnosis and management of methylmalonic and propionic acidemia. Orphanet J Rare Dis 9, 130 (2014).

42. Chandler, R.J. et al. Mitochondrial dysfunction in mut methylmalonic acidemia. FASEB J 23, 1252–61 (2009).

43. Manoli, I. et al. FGF21 underlies a hormetic response to metabolic stress in methylmalonic acidemia. JCI Insight 3 (2018).

44. Wilnai, Y., Enns, G.M., Niemi, A.K., Higgins, J. & Vogel, H. Abnormal hepatocellular mitochondria in methylmalonic acidemia. Ultrastruct Pathol 38, 309–14 (2014).

45. Zsengeller, Z.K. et al. Methylmalonic acidemia: a megamitochondrial disorder affecting the kidney. Pediatr Nephrol 29, 2139–46 (2014).

46. Molema, F. et al. Fibroblast growth factor 21 as a biomarker for long-term complications in organic acidemias. J Inherit Metab Dis 41, 1179–1187 (2018).

47. Suomalainen, A. et al. FGF-21 as a biomarker for muscle-manifesting mitochondrial respiratory chain deficiencies: a diagnostic study. Lancet Neurol 10, 806–18 (2011).

48. Inagaki, T. et al. Endocrine regulation of the fasting response by PPARalpha-mediated induction of fibroblast growth factor 21. Cell Metab 5, 415–25 (2007).

49. Lundasen, T. et al. PPARalpha is a key regulator of hepatic FGF21. Biochem Biophys Res Commun 360, 437–40 (2007).

50. Ozaki, Y. et al. Rapid increase in fibroblast growth factor 21 in protein malnutrition and its impact on growth and lipid metabolism - ERRATUM. Br J Nutr 114, 1535–6 (2015).

51. Laeger, T. et al. FGF21 is an endocrine signal of protein restriction. J Clin Invest 124, 3913–22 (2014).

52. von Holstein-Rathlou, S. et al. FGF21 Mediates Endocrine Control of Simple Sugar Intake and Sweet Taste Preference by the Liver. Cell Metab 23, 335–43 (2016).

53. Ables, G.P., Perrone, C.E., Orentreich, D. & Orentreich, N. Methionine-restricted C57BL/6J mice are resistant to diet-induced obesity and insulin resistance but have low bone density. PLoS One 7, e51357 (2012).

54. Hotta, Y. et al. Fibroblast growth factor 21 regulates lipolysis in white adipose tissue but is not required for ketogenesis and triglyceride clearance in liver. Endocrinology 150, 4625–33 (2009).

55. Owen, B.M. et al. FGF21 acts centrally to induce sympathetic nerve activity, energy expenditure, and weight loss. Cell Metab 20, 670–7 (2014).

56. Kliewer, S.A. & Mangelsdorf, D.J. A Dozen Years of Discovery: Insights into the Physiology and Pharmacology of FGF21. Cell Metab 29, 246–253 (2019).

57. Kim, K.H. et al. Autophagy deficiency leads to protection from obesity and insulin resistance by inducing Fgf21 as a mitokine. Nat Med 19, 83–92 (2013).

58. Vernochet, C. et al. Adipose tissue mitochondrial dysfunction triggers a lipodystrophic syndrome with insulin resistance, hepatosteatosis, and cardiovascular complications. FASEB J 28, 4408–19 (2014).

59. Shimizu, I. et al. Vascular rarefaction mediates whitening of brown fat in obesity. J Clin Invest 124, 2099–112 (2014).

60. Badman, M.K. et al. Hepatic fibroblast growth factor 21 is regulated by PPARalpha and is a key mediator of hepatic lipid metabolism in ketotic states. Cell Metab 5, 426–37 (2007).

61. Pawlak, M., Lefebvre, P. & Staels, B. Molecular mechanism of PPARalpha action and its impact on lipid metabolism, inflammation and fibrosis in non-alcoholic fatty liver disease. J Hepatol 62, 720–33 (2015).

62. Xu, J. et al. Peroxisome proliferator-activated receptor alpha (PPARalpha) influences substrate utilization for hepatic glucose production. J Biol Chem 277, 50237–44 (2002).

63. Dewulf, J.P. et al. The synthesis of branched-chain fatty acids is limited by enzymatic decarboxylation of ethyl-and methylmalonyl-CoA. Biochem J 476, 2427–2447 (2019).

64. Rozman, J., Klingenspor, M. & Hrabe de Angelis, M. A review of standardized metabolic phenotyping of animal models. Mamm Genome 25, 497–507 (2014).

65. Rekant, S.I., Lyons, M.A., Pacheco, J.M., Arzt, J. & Rodriguez, L.L. Veterinary applications of infrared thermography. Am J Vet Res 77, 98–107 (2016).

66. de Menezes, R.C., Ootsuka, Y. & Blessing, W.W. Sympathetic cutaneous vasomotor alerting responses (SCVARs) are associated with hippocampal theta rhythm in nonmoving conscious rats. Brain Res 1298, 123–30 (2009).

67. Saegusa, Y. & Tabata, H. Usefulness of infrared thermometry in determining body temperature in mice. J Vet Med Sci 65, 1365–7 (2003).

68. Johnson, S.R., Rao, S., Hussey, S.B., Morley, P.S. & Traub-Dargatz, J.L. Thermographic Eye Temperature as an Index to Body Temperature in Ponies. Journal of Equine Veterinary Science 31, 63–66 (2011).

69. Sanchez-Alavez, M., Alboni, S. & Conti, B. Sex- and age-specific differences in core body temperature of C57Bl/6 mice. Age (Dordr) 33, 89–99 (2011).

70. Lecorps, B., Rodel, H.G. & Feron, C. Assessment of anxiety in open field and elevated plus maze using infrared thermography. Physiol Behav 157, 209–16 (2016).

71. Speakman, J.R. Measuring energy metabolism in the mouse - theoretical, practical, and analytical considerations. Front Physiol 4, 34 (2013).

72. Sundberg, J.P. et al. Primary follicular dystrophy with scarring dermatitis in C57BL/6 mouse substrains resembles central centrifugal cicatricial alopecia in humans. Vet Pathol 48, 513–24 (2011).

73. Rathkolb, B. et al. Clinical Chemistry and Other Laboratory Tests on Mouse Plasma or Serum. Curr Protoc Mouse Biol 3, 69–100 (2013).

