## Supplementeray Information for "Insights into energy balance dysregulation from a mouse model of methylmalonic aciduria"

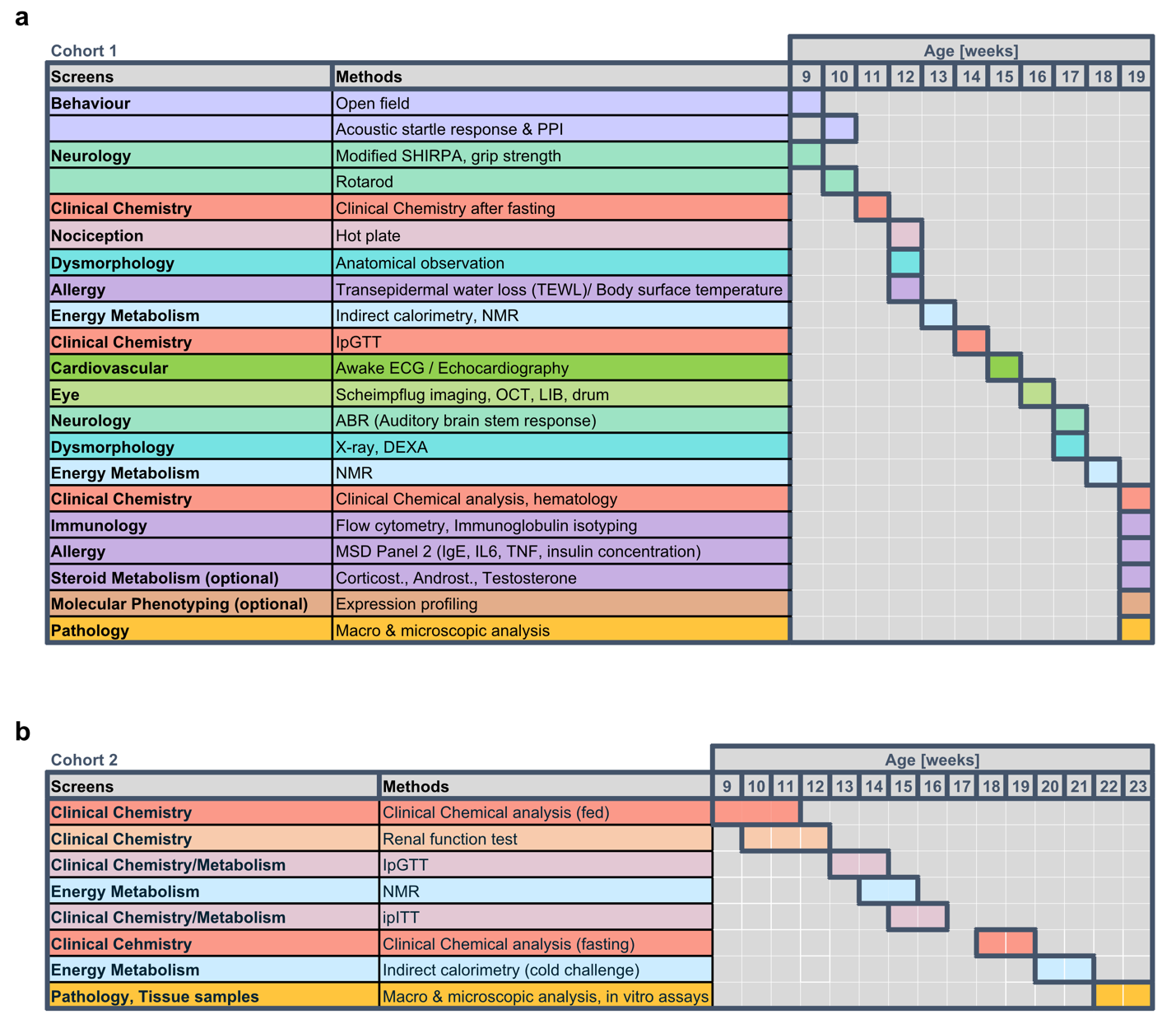

**Supplementary Figure 1. Experimental workflow of the 2 mice cohorts. a** Mice in the first cohort were examined in a an experimental workflow modified after Gailus-Durner 2009, Lucienne 2020). **b** Mice in the second cohort were examined in the depicted bespoke experimental workflow.

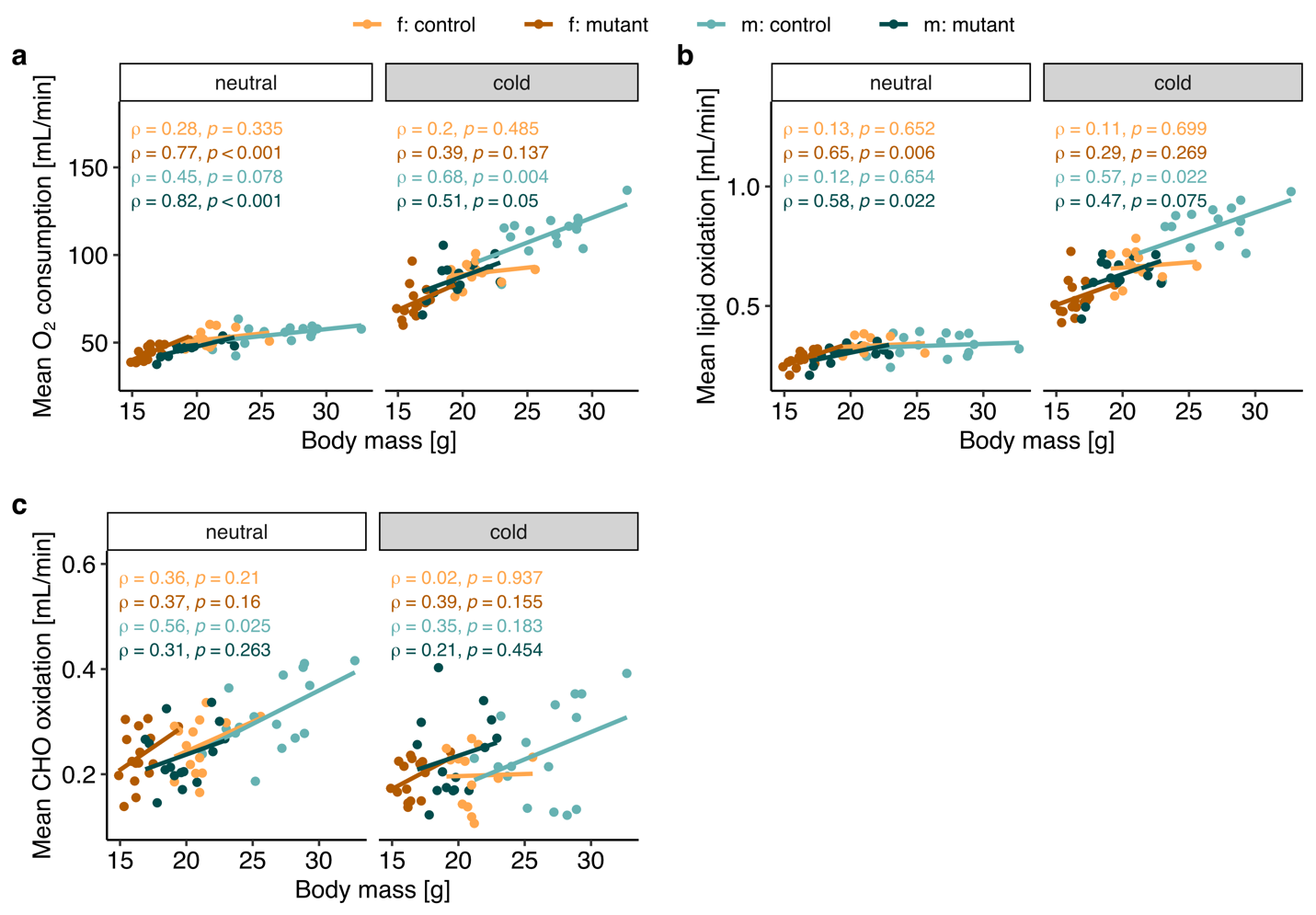

**Supplementary Figure 2. Linear models of indirect calorimetry before and during cold challenge.** **a** Oxygen consumption, **b** lipid oxidation, and **c** carbohydrate (CHO) oxidation of single mouse from Figure 3 separated according to neutral or cold environmental condition.

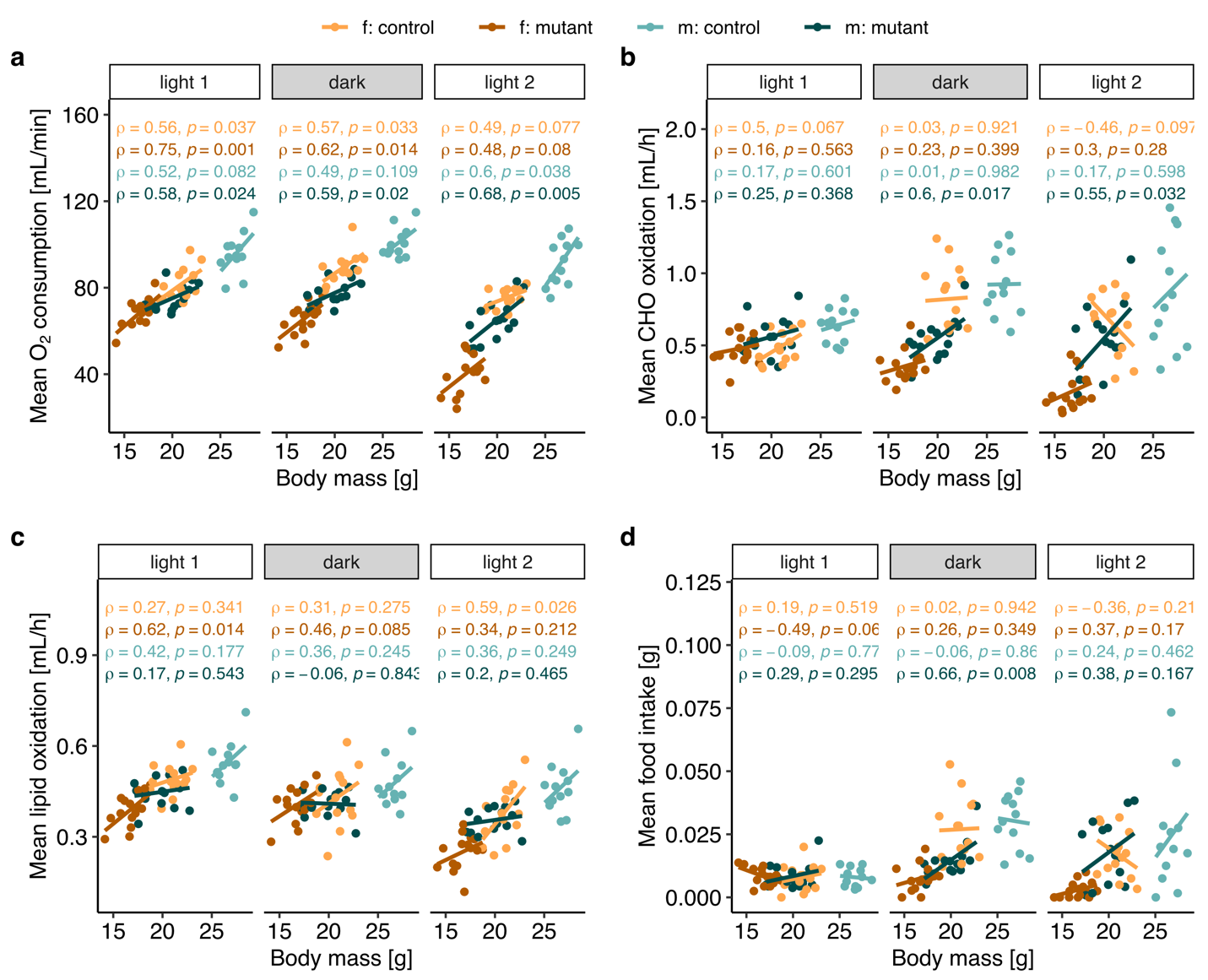

**Supplementary Figure 3. Linear models of indirect calorimetry in standard conditions.** **a** Oxygen consumption, **b** lipid oxidation, and **c** carbohydrate (CHO) oxidation. of single mouse from Figure 4 according to light and dark phases.

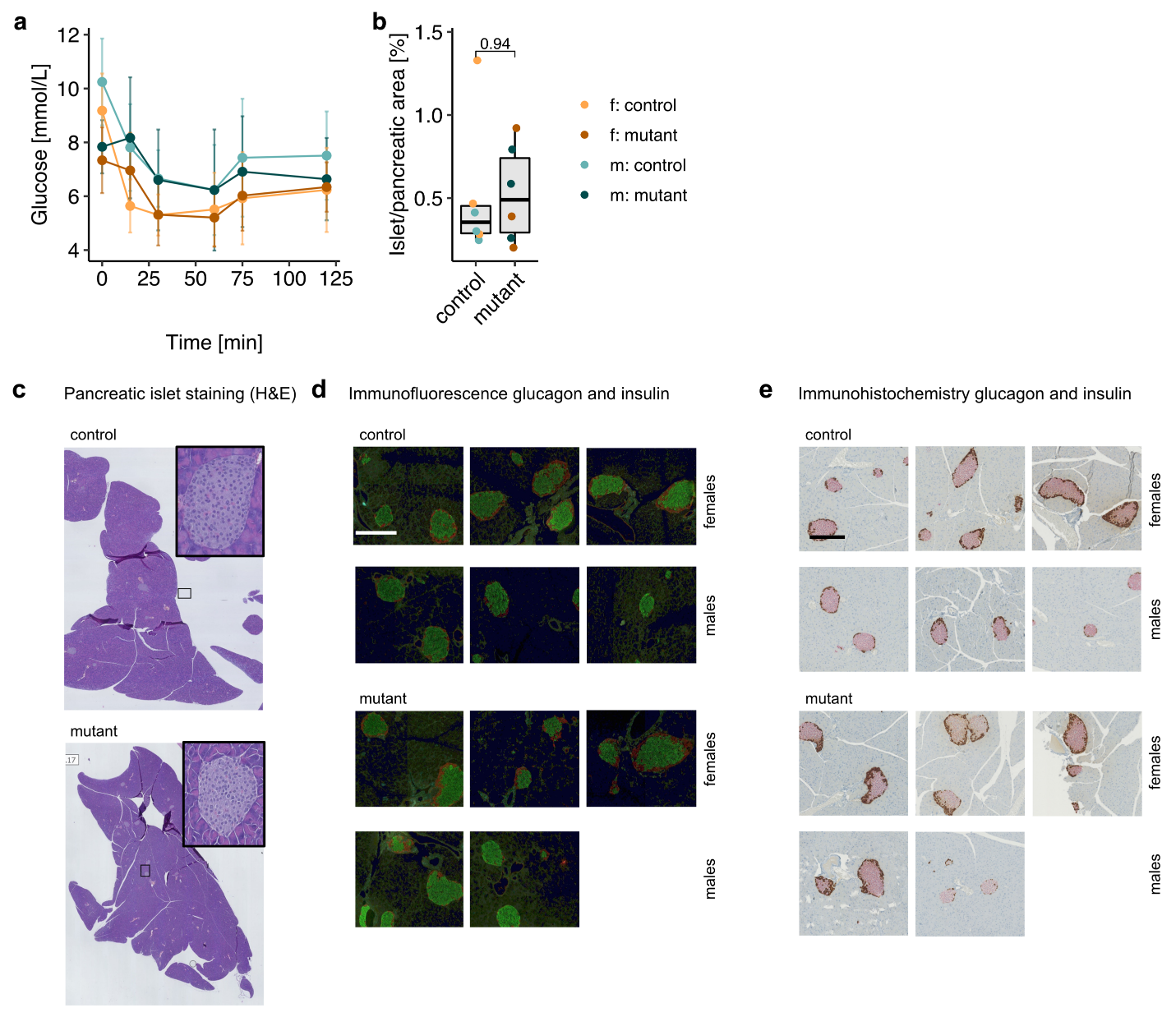

**Supplementary Figure 4. a** Glucose excursion after intraperitoneal injection of insulin. n=15 *Mmut*-ki/wt females, n=16 *Mmut*-ko/ki females, n=17 *Mmut*-ki/wt males, n=16 *Mmut*-ko/ki males, 16 weeks of age. Mean +/- SD is shown. **b** Ratio of islet area compared to pancreatic area determined from n=3 *Mmut*-ki/wt females, n=3 *Mmut*-ko/ki females, n=2 *Mmut*-ki/wt males, n=2 *Mmut*-ko/ki males. p-value determined by Wilcoxon test. **c** Representative H&E of pancreas including islet (inset) of a control and mutant mouse. **d-e** Representative islet immunohistochemistry of glucagon and insulin detected by **d** immunofluorescence (insulin: green, glucagon: red) and **e** chromogenic labelling (insulin: dark red, glucagon: light red). Each panel represents a different mouse. Scale bar represents 250 µm. (f. females, m. males, mutant: *Mmut-ko/ki*, control: *Mmut*-ki/wt)

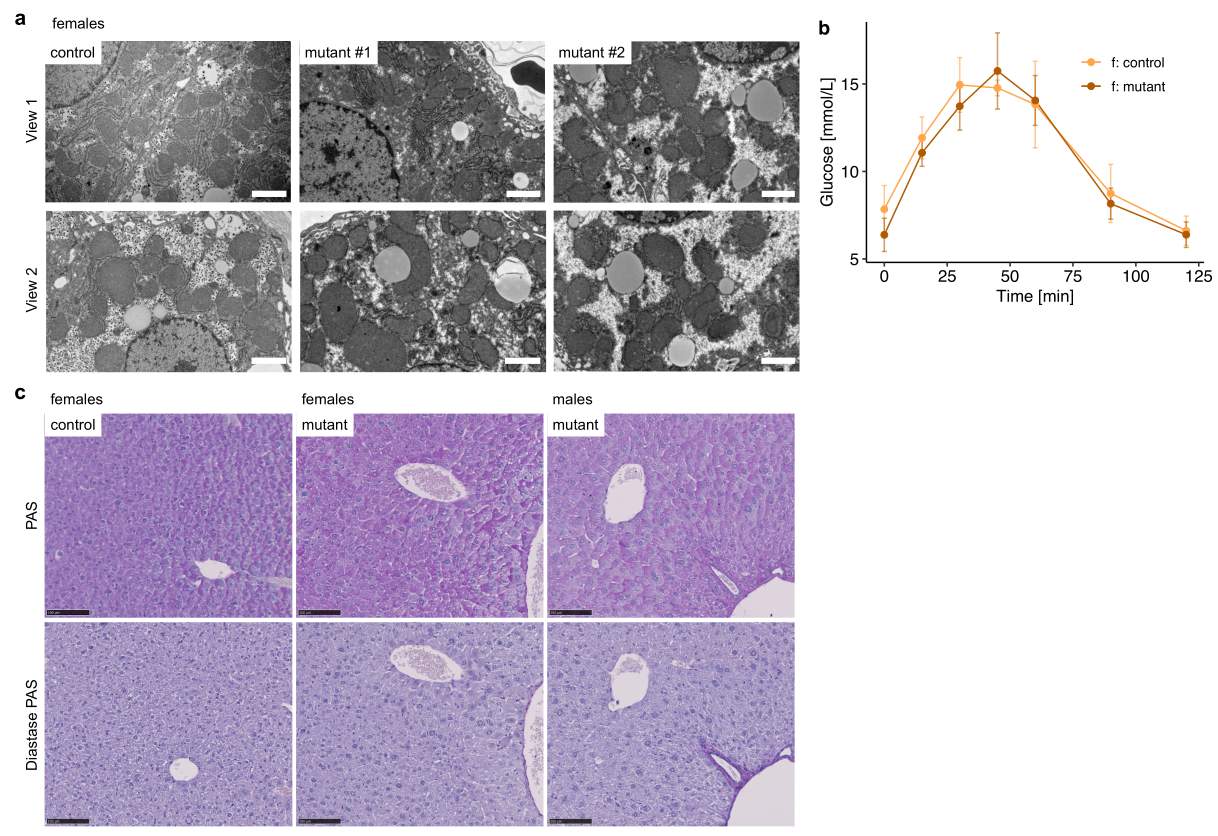

**Supplementary Figure 5. a** Transmission electron microscope images of liver samples from two distinct *Mmut*-ko/ki mice and a *Mmut*-ki/wt mouse, each 7 months of age. Scale bar represents 2 µm. **b** Glucose excursion after intraperitoneal injection of pyruvate. n=7 *Mmut*-ko/ki females and n=7 *Mmut*-ki/wt females mice, 5.5 months of age. Mean +/- SD is shown. **c** Upper panel: serial liver sections stained with PAS (periodic acid-Schiff) to detect polysaccharides such as glycogen. Magenta color represents PAS positive. Lower panel: PAS staining of the same sections following digestion with diastase (which must be PAS negative). Scale bar represents 100 µm.

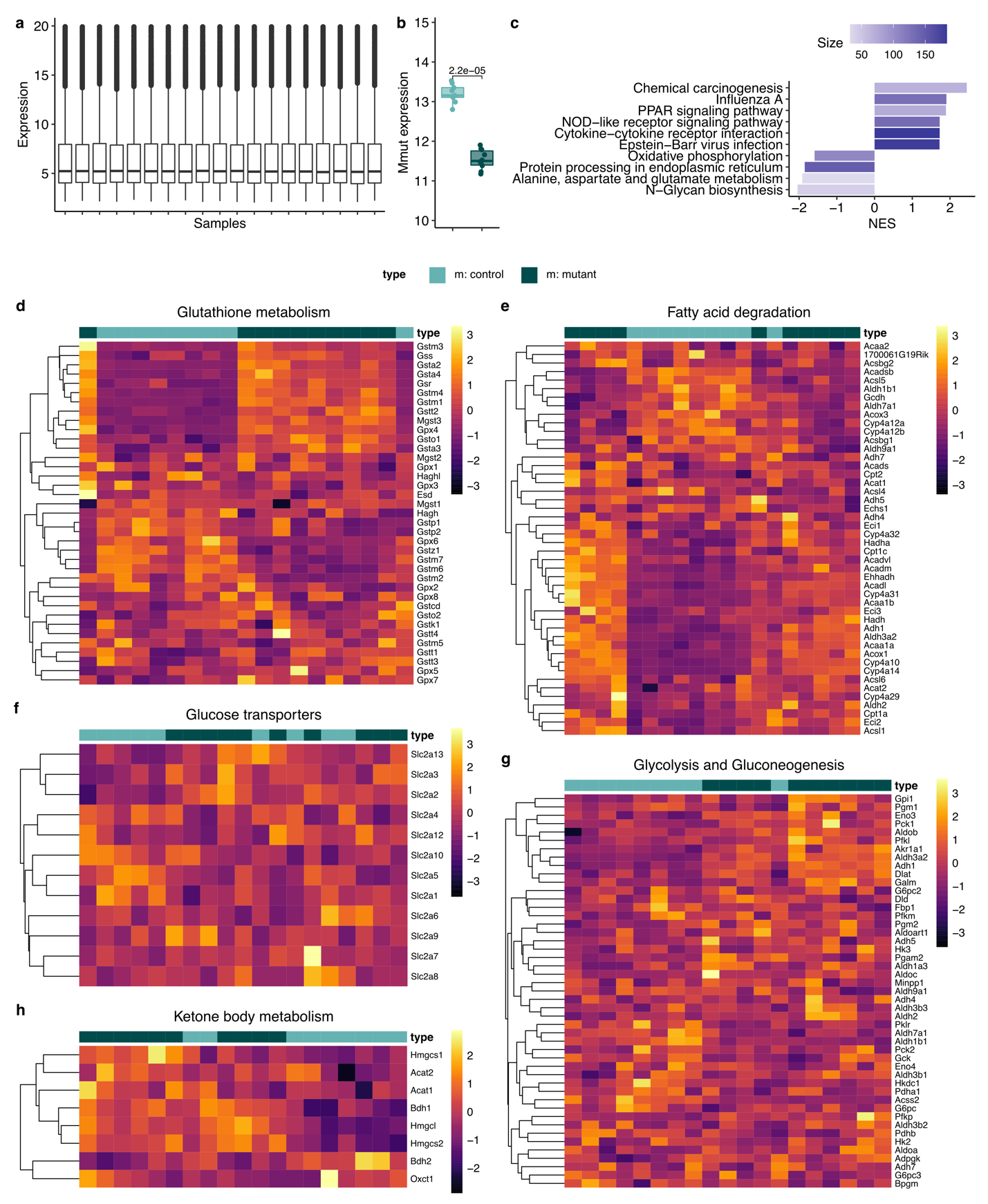
**Supplementary Figure 6.** **a** Total microarray signal for each sample. **b** Expression of *Mmut* detected by microarray; p value calculated by Wilcoxon test. **c** KEGG gene-set enrichment analysis. **d-h** Heat maps of all genes according to KEGG belonging to **d** glutathione metabolism, **e** fatty acid degradation, **f** glucose transport, **g** glycolysis and gluconeogenesis, and **h** ketone body metabolism. All samples derived from RNA from liver tissue of 9 *Mmut*-ki/wt and 10 *Mmut*-ko/ki male mice in the *ad libitu*m fed state.

| **Genes** | **Primer sequences** |
| --- | --- |
| *Acaa1b* | F: 5‘-AGA CAT CTC CGT GGG CAA TG-3‘  R: 5‘-CTG CAG TCC CGA TGA ACA CT-3‘ |
| *Acadl* | F: 5‘-GGT GTT CAT CAC TAA TGG CTG G-3‘  R: 5‘-AGT TCT GCT GTG TCC TGA GC-3‘ |
| *Acadm* | F: 5‘-TGA CAA AAG CGG GGA GTA CC-3‘  R: 5‘-TTT CCG GAA TGT GCG CGT TG-3‘ |
| *Actb* | F: 5‘-GGT GGG AAT GGG TCA GAA GG-3‘  R: 5‘-AGG TCT CAA ACA TGA TCT GGG T-3‘ |
| *Cyp4a10* | F: 5‘-GAC CTA CCT CCA GGC CAT TG-3‘  R: 5‘-AGC TTG ATC ACT CCA TCT GTG T-3‘ |
| *Cyp4a31* | F: 5‘-CCG GAA GAT GCT AAC CCC AG-3‘  R: 5‘-GCC GTT CCC ATT TGT CTA GC-3‘ |
| *Ehhadh* | F: 5‘-ACA ACT TCT GTG CAG GTG CT-3‘  R: 5‘-GAA GCC AAC ACG AGC CTT TG-3‘ |
| *Fgf21* | F: 5‘-GCT CTC TAT GGA TCG CCT CAC-3‘  R: 5‘-GAG TCA GGA CGC ATA GCT GG-3‘ |
| *Hmgcl* | F: 5‘-AGG CTT TGA GGA AGC GGT AG-3‘  R: 5‘-CTT TAG CCG GGG AGA CCT TC-3‘ |
| *Hmgcs1* | F: 5‘-TCT ACC GCA AAA AGA TCC GTG-3‘  R: 5‘-TCT AAT TTA ACG TCC CCA AAG GC-3‘ |
| *Hnf4a* | F: 5‘-CTG TCC CAG CAG ATC ACC TC-3‘  R: 5‘-TCA TTG CCT AGG AGC AGC AC-3‘ |
| *Pck1* | F: 5‘-TGC ATG AAA GGC CGC ACC-3‘  R: 5‘-GTT GCA GGC CCA GTT GTT G-3‘ |
| *Pklr* | F: 5‘-GGG TGA CCT TGG CAT TGA GA-3‘  R: 5‘-TGC TCT CCA GCA TCT GTG TG-3‘ |
| *Ppargc1a* | F: 5‘-CCT CAC ACC AAA CCC ACA GA-3‘  R: 5‘-GAG GAG TTA GGC CTG CAG TT-3‘ |

**Supplementary Table 1: List of primer sequences used for qRT-PCR analysis.** F: forward primer. R: reverse primer
